# Kat5 cKO Biological Domain Signatures Align with Human Alzheimer’s Disease

**DOI:** 10.1101/2025.09.13.676046

**Authors:** GA Cary, JE Young, SE Rose, H Frankowski, M Darvas, M Bothwell, S Jayadev, AN Reid, A Greenwood, A Levey, K Leal, GW Carter, JC Wiley

**Affiliations:** University of Kansas, 1450 Jayhawk Blvd, Lawrence, KS 66045; University of Washington School of Medicine, 1959 NE Pacific St, Seattle, WA 98195; The Jackson Laboratory, 600 Main St Bar Harbor, ME 04609; Emory University School of Medicine, 100 Woodruff Circle. Atlanta, GA 30322 USA; Sage Bionetworks, 2901 3rd Ave #330, Seattle, WA 98121

**Keywords:** Alzheimer’s Disease, Genetics, Transcriptomics, Proteomics, Biological Domain, Endophenotype, Epigenetics, Chromatin

## Abstract

**BACKGROUND:** Alzheimer’s disease (AD) is associated with amyloid plaques and can be caused by autosomal dominant mutations in APP or PSEN1/2, which form an enzyme substrate complex. Decreases in catalysis of AD mutant APP and PSEN1 supports the hypothesis that membrane delimitation of KAT5 could contribute to AD.

**METHODS:** We compare the hippocampal transcriptome profiles of the Kat5 brain-specific knockout mouse to multiple AD datasets through alignment with the TREAT-AD AD biological domains. We examine KAT5 subcellular localization in human WT and AD neurons.

**RESULTS:** The Kat5 KO mouse demonstrates downregulation of synaptic genes, metabolic pathways, and upregulation of DNA replication and repair, cell cycle and immune response genes. We see similar profiles in Kat5 and comparative AD datasets. KAT5 is restricted to the cytosol in human AD neurons.

**DISCUSSION:** This analysis supports the hypothesis that KAT5 nuclear signaling down stream of APP cleavage plays a pivotal role in neuronal homeostasis and immune regulation.

## 1 Background

The cloning of the β-amyloid precursor protein (APP) sparked speculation about its potential role as a receptor, given its large extracellular domain and small intracellular domain that interacts with multiple signaling factors(1). However, the lack of an identified ligand or defined signaling mechanism cast doubt on this hypothesis, leaving APP’s biological function unclear. APP is a type-I single-pass transmembrane protein sequentially cleaved in its extracellular domain by either α- or β-secretase, followed by cleavage within the transmembrane region by the heterotetrameric γ-secretase complex. This process begins at the ε-site near the cytosol and proceeds in three-amino-acid increments to the γ-sites, producing amyloid peptides(2, 3).

APP is crucial for development, as genetic deletion of two out of the three APP isoforms leads to early postnatal lethality(4, 5). Numerous studies suggest that APP plays a role in neurodevelopment, including synapse formation, axon pathfinding, and synaptic plasticity within developing cortical layers(6–10). Supporting this, cultured neurons derived from APP-null mice, while retaining amyloid beta precursor like protein 1 and 2 (APLP1 and APLP2), show a reduced number of synapses(11). Furthermore, the brain specific triple knockout of all three isoforms of the APP family did not result in embryonic lethality, but did cause profound alterations in synaptic signaling and plasticity along the performant path fibers, strongly indicating APP’s role in synaptic function(12).

Mutations in APP near the β- and γ-secretase cleavage sites lead to autosomal dominant Alzheimer’s disease(1). The γ-secretase complex, with presenilin (PSEN1 or PSEN2) at its catalytic core, directly associates with APP and mediates its intramembranous aspartyl-dependent proteolysis(2). Mutations in PSEN1 or PSEN2 also cause autosomal dominant AD(1, 13). While amyloid production has been the key focus in explaining the link between these mutations and AD, many studies support an alternative “loss of function” hypothesis. These reports show that familial AD (FAD) mutations often reduce overall proteolytic activity and sometimes decrease levels of major amyloid species(14–18). Additionally, some FAD mutations do not increase Aβ42 production but instead lower Aβ40 levels or produce longer Aβ peptides predicted to remain embedded in the membrane(19, 20). Recently, a study demonstrated that FAD mutations in APP and PSEN1 stall the Presenilin-APP enzyme-substrate complex, decrease AICD production, and support the plausibility that FAD mutations could lead to a disruption of APP cleavage-dependent signaling(21).

Over two decades ago, the association between APP and APBB1/FE65 and KAT5/TIP60 was identified and implicated as a Notch-like signaling mechanism(22, 23). Disputes over the exact manner of transcriptional regulation, and the genes regulated, cast some doubt upon this signaling modality(22, 24–26). KAT5 is a major histone acetylase in brain, and accordingly, epigenetic regulation of chromatin accessibility may be impacted by KAT5 release from the membrane. Elevated histone deacetylation is implicated in AD(27, 28), and as KAT5 operates in opposition to histone deacetylases (HDACs), a decrement in histone acetylase activity could promote hypoacetylation, decreasing epigenetic regulation involved in synaptic plasticity(29, 30).

In this study, we propose that decreased γ-secretase function, likely due to a stalled transition-state intermediate of presenilin and APP, causes KAT5 to become sequestered at the membrane, repressing nuclear signaling. If APP/KAT5 nuclear signaling plays a role in AD pathogenesis, we would expect to see similar transcriptomic signatures in the KAT5 brain-specific inducible knockout and AD associated transcriptomic profiles. The brain-specific inducible Kat5 knockout mouse (Kat5 cKO) was generated by Gregor Eichle’s lab as previously reported(31). Here we report that deletion of KAT5 results in a loss of synaptic gene expression, increases in cell cycle regulation, and induction of immune function partially mirroring patterns in four distinct AD transcriptomic datasets (5XFAD, LOAD, EOAD, and FAD). We further demonstrate that KAT5 is restricted from the nucleus specifically in human AD neurons and is tightly coupled with neuronal survival, likely mediated through the downregulation of genesets negatively associated with neurodegeneration (aka, synaptic genes). These results support a plausible role for disrupted KAT5 nuclear signaling in AD pathogenesis.

## 2 Methods

### 2.1 Datasets and data availability

#### 2.1.1 Kat5 cKO mouse model data

The Kat5 mouse RNAseq model data was extracted from the published report from the Eichle laboratory(31). The dataset contained the complete gene expression profile observed within the Kat5 forebrain-specific tamoxifen-inducible CamKIIα promoter driven CRE mouse crossed onto the floxed Kat5 mouse. The CA1 region of the hippocampus was laser dissected after 10 days of tamoxifen induction of the CRE-recombinase within 2-month-old mice. The dataset included 6 experimental (CamKII-CRE X Kat5_flox_) and 6 controls (Kat5_flox_) mice, both treated with tamoxifen to control for non-specific drug treatment effects. The fold-change calculations were made across all genes, and statistical significance was determined using an adjusted p-value threshold of 0.05 independent of fold-change level. The dataset is parsed into low (1.2 fold), medium (1.5 fold) and high (2 fold) levels of up or downregulation. The analysis performed assessing global shifts in transcriptomic signatures employ the 1.2-fold change group, to attain the highest level of biological process sensitivity.

#### 2.1.2 Human Early-Onset AD and Familial AD datasets

The early onset AD and FAD human datasets are extracted from Antonell et al(32) in which sporadic EOAD patients and FAD patients were interrogated post-mortem to compare the transcriptomic profiles of these two types of AD. Seven subjects were employed in the sporadic EOAD (under 65 years of age), Presenilin-1 FAD (M139T), and the control group, with a preponderance of males in all groups: 5/7, 6/7, and 4/7 respectively. The average age of the participants varies from 50 for the controls, 54 for the FAD participants, and 63 for the sporadic AD patients. The APOE allele was predominantly APOE3 across subject groups with a single e3/e4 heterozygous participant in each study group. The neuropathological rating was performed using the ABC scale (A: Thal Amyloid rating, B: Braak Tau stage, C: CERAD amyloid presence), and the sporadic EOAD and FAD subjects rated severe (A3, B3 and C3) across pathology measures —the controls had no noted pathology. RNA was extracted from the posterior cingulate cortex, purified and quality control assessed with a high value RNA integrity number (RIN) across samples, average 7.3 for experimental groups and 7.2 for control participants. The labeled RNA was hybridized to Affymetrix ST 1.1 Human Gene chip, the data was cleaned and compared between control and experimental groups using Bioconductor R packages developed for Affymetrix analysis and LIMMA packages(32). The differential gene expression was identified using pairwise comparisons between experimental and control groups and corrected for false discovery using Benjamini and Hochberg methods. The data can be accessed at Gene Expression Omnibus (GEO) accession GSE39420. For additional details, see the original characterization(32).

#### 2.1.3 Human Late Onset AD Dataset

The late onset AD (LOAD) dataset was developed from multiple brain banks representing large independent research efforts within AMP-AD and TREAT-AD. We pulled the transcriptomic datasets together within our bioinformatics pipeline analysis to facilitate TREAT-AD target evaluation and grouping into the biological domains developed to represent the AD endophenotypes in an objective fashion(33). The compiled datasets are currently available on the AD Knowledge Portal(33). The transcriptomic dataset was drawn from ROS-MAP, Mayo, and Mount Sinai brain banks, comprising over 1700 individual brains across genders, ethnicities and age brackets. No subordinate evaluation of the subpopulations within the collective dataset was initially performed, and hence only the collective dataset was analyzed and compared to the Kat5 cKO model in the present work.

#### 2.1.4 The 5XFAD Mouse

The 5XFAD transgenic mouse strain carries 5 distinct APP and PSEN1 FAD mutations that drive elevated amyloid production and memory deficits early in mouse development(34). The 5X FAD mouse transcriptome is employed and rigorously evaluated in the MODEL-AD consortia efforts(35–37). The transcriptomic dataset for the 12-month 5XFAD mouse model was employed in the comparison to the Kat5 cKO dataset. The transcriptomic datasets for the 5XFAD mouse are available on the AD Knowledge Portal (https://adknowledgeportal.synapse.org; synapse ID: syn66318332).

#### 2.1.5 Human Frontotemporal Dementia Dataset

We examined the transcriptomic data emerging from the Risk and modifying factors in Frontotemporal Dementia (RiMod-FTD) consortium(38), specifically comparing the differential expression in the case-control study of FTD patients with MAPT mutations to the Kat5 KO mouse model differential expression profile to assess AD specificity of the Kat5 molecular signature alignment.

### 2.2 GSEA Analysis of the Biological Domains

#### 2.2.1 Biological domain enrichment

All datasets were aligned to the TREAT-AD biological domains employing gene set enrichment analysis (GSEA) to the annotated GO terms across the three tiers of GO. The enriched GO terms were organized into the biological domains based on the definition profiles of the AD biological domains(33) and the biological domains were ranked based on the frequency of enriched term participation. To evaluate the comparative enrichment of various biological domains and their respective Gene Ontology (GO) terms, gene set enrichment analysis (GSEA) was conducted utilizing the gseGO function within the clusterProfiler R package(39). Subsequently, the outcomes were grouped into biological domains according to the enriched terms’ GO ID. Each enrichment analysis was conducted using non-zero log-fold change values, sorted in descending order. The presented outcomes comprise the normalized enrichment score (NES) and the Benjamini-Hochberg corrected p-value (p adj) for GO terms associated with each biological domain.

#### 2.2.2 Shared biological domain GO term enrichment between models

All pairwise comparisons made between the Kat5 cKO and the AD datasets were enriched for the GO terms mapping onto the 19 biological domains. The GO terms shared between the groups were mapped per biological domain, the domains with shared patterns of enrichment were identified and presented. The enrichment draws from the transcriptomic datasets for each model and accordingly contains directional information about change in regulation. Only domains with at least five terms of coordinate regulation are considered significantly overlapping and surfaced in the present work.

#### 2.2.3 Correlation Analysis Between Datasets

The gene correlation analysis between the Kat5 cKO differentially expressed genes and those differentially expressed in AD models employ Pearson correlation performed by R stat package in BioConductor. The Pearson r correlation term shows the overlap between the whole DEG set models, without stratification into biological domains or GO terms. The p-value calculation is performed to assess the statistical relevance of the associated correlation (r) value and the beta coefficient index provides the interaction slope of the compared models.

### 2.3 Kat5 cKO DEG Kappa-Network Modeling

The kappa-network approach identifies GO biological process (BP) enrichments based upon the DEGs. These networks form an exhaustive set of related processes, linked by genes annotated to multiple GO terms, and the degree of term conservation is the linking edge between nodes. These are mapped based on their derivation from either up- or downregulated DEGs, so process families can be explored simultaneously based upon (1) significance level, (2) degree of conservation of genes between terms (kappa-value), and (3) the directionality of DEG shift. The network modeling was performed using the cytoscape application Clue-GO with a kappa-value set at 0.5, and a selectivity for specific directionality of 0.6 for inclusion into either the downward or upward directed network. The grouping of terms observed in Figure 1 was based upon the best term characterization of the boxed subcomponent of the network observed in either regulation direction. The calculation of the total fraction of top-level terms was tallied and grouped based upon the most enriched term being lead with ranking elevation given to the highest ontological position within GO within the cluster.

**Figure 1.**
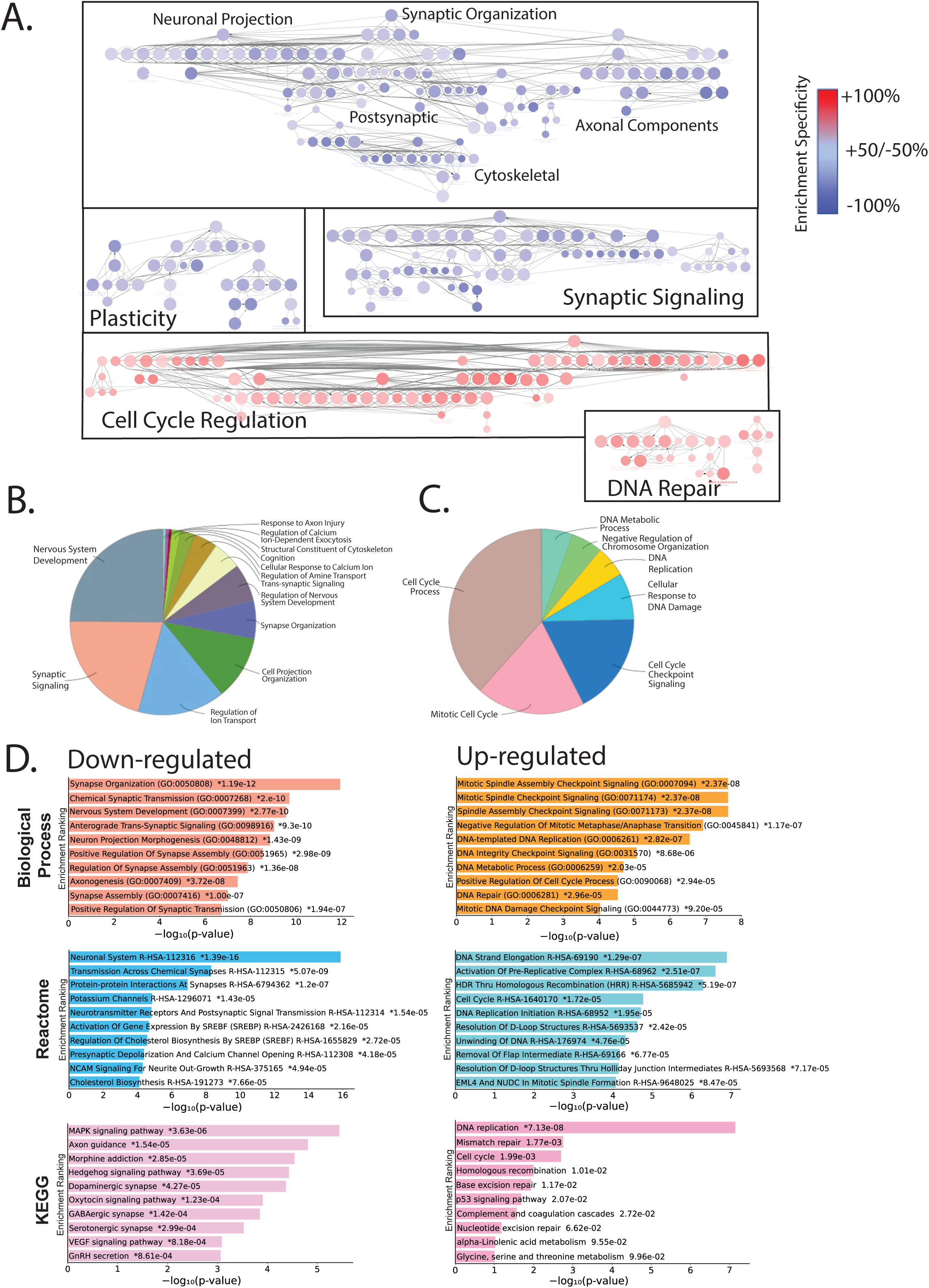
Up and Downregulated Processes in the KAT5 cKO. (A) Kappa-networks were constructed from the up- and downregulated gene sets derived from the KAT5 cKO mouse CA1 hippocampus. The nodes represent Gene Ontology (GO) biological process terms, and the linking edges represent the kappa-score derived from shared genes annotated between GO terms. The annotation of the subclusters of the network represent the broader biological families of impacted biological processes. The blue nodes are downregulated, and are predominantly related to synaptic linked processes, with development, plasticity and synaptic signaling being the three largest families.Upregulated processes are shown in red and are predominantly centered upon DNA metabolic and cell cycle functions. (B) All the downregulated processes are represented by a pie chart showing the major themes and fractional composition of the GO term annotation, in which synaptic development and function are the predominant fractions. (C) The upregulated terms are represented by fractional contribution of enrichment, demonstrating specific classes of DNA metabolic and cell cycle events playing the largest role. The top ten terms from each branch of GO BP (D, top), KEGG pathway analysis (D, middle), and Reactome pathway mapping (D, bottom) are shown, with the downregulated processes on the left and the upregulated processes on the right.

### 2.4 The characterization of enrichment of Kat5 DEGs

The enrichment of terms within each branch of the GO employed the R based web-accessible tool Enrichr (https://maayanlab.cloud/Enrichr/)(40). Here the top 10 terms within the biological process branch of the GO were mapped for both the upward and downward regulated gene sets, to provide a direct comparison to the kappa-network enrichment. Similarly, the top ten associated KEGG and Reactome pathways were also demonstrated for the up and downregulated gene sets. The upregulated and downregulated DEGs within the Kat5 cKO model at each tier (1.2, 1.5, and 2-fold) were compared to the 2023 GWAS Catalogue (which incorporates both TopMed and UK Biobanks data)(41) to assess enrichment of genetic trait linkage using the Enrichr tool(42).

### 2.3 iPSC Neuronal models employed

#### 2.3.1 iPSC Neuronal lines and culture conditions

Human induced pluripotent stem cells (hiPSCs) harboring the APP Swedish (KM670/671NL), Presenilin-2 N141I or an APP duplication were differentiated into excitatory cortical neurons following published protocols(43, 44). Control cells were either isogenic WT hiPSCs (for APP Swedish) or cells derived from a non-demented subject (for APP Duplication). Neuronal cultures were enriched by depletion of non-neuronal cell types using antibodies for CD184PE and CD44PE as previously described(45). Following neuronal purification, cells were replated for five days on Geltrex coated coverslips prior to immunocytochemistry, and when relevant neurons were treated with 1uM DAPT (Sigma) 24 hours prior fixation. APP Swedish hiPSCs and APP duplication hiPSCs have been characterized and published(46, 47). PSEN2 N141I variant was introduced into control cells using CRISPR-Cas9 genome engineering following established protocols(44).

#### 2.3.2 KAT5 immunofluorescent imaging and analysis in hiPSC-derived neurons

Imaging was performed on a Leica TCS SP8 confocal laser microscope with a 20x lens. Images were acquired in the Leica Application Suite X (LAS X 3.5.519976) with the following settings: 405 nm laser at 1% and HyD detector 1 gain at 100%; 488 nm laser at 1% and PMT detector gain 1247 V; 552 laser at 1% and HyD detector 2 gain at 100%. Image format was 1024 x 1024, speed 400 Hz, and 7x zoom. Image analysis was performed on 5 neurons per genotype using Bitplane Imaris software (Oxford Instruments). A surface object workflow was used in Imaris to define the DAPI+ nuclear ROI. Neuronal nuclei were identified by the presence of MAP2 cytoplasmic staining. The nuclear ROI surface was used to mask the KAT5 (green) channel and isolate nuclear KAT5 staining. A surface object workflow was run on the masked KAT5 channel to quantify nuclear KAT5 fluorescent intensity in the neurons.

A one-way ANOVA and Holm-Šídák’s multiple comparisons test were used to compare nuclear KAT5 fluorescent intensities between the familial AD hiPSC-derived neurons and wildtype neurons. A Mann-Whitney test was used to compare nuclear KAT5 fluorescent intensities between the DMSO vehicle control and DAPT-treated wildtype hiPSC-derived neurons.

## 3. Results

The goal of this work is to assess the plausibility of disruption to KAT5 nuclear chromatin regulation as a driving component of AD pathogenesis by comparing the Kat5 cKO transcriptomic signature(31) to murine and human AD datasets to assess their similarity. First, we explore the enrichment signatures of the Kat5 cKO mouse using complementary techniques of kappa-network models and gene set enrichment analysis (GSEA). The kappa-network methodology is employed to capture the greatest breadth of altered processes within the KAT5 model, while the GSEA approach focuses on gene ontology (GO) term and pathway specific enrichment patterns. The enrichment of the downregulated gene set highlighted predominantly synaptic genes (Figure 1A blue, B) in keeping with the original characterization(31). The synaptic gene downregulation network clusters into three subnetworks that align with synaptic developmental, plasticity, and synaptic signaling. The upregulated kappa-network cluster into cell cycle and DNA repair subnetworks (Figure 1A red, C).

The enrichment of Kat5 DEGs shows down-regulated processes and pathways in the left column and up-regulated on the right (Figure 1D). The top 10 down-regulated GO BPs are all synaptic, split between structural, developmental and signaling related terms, while up-regulated BPs are cell cycle and DNA replication and repair focused, consistent with the kappa-network analysis. These results are mirrored in both the Reactome and KEGG pathway enrichments (second and third rows, Figure 1D), with Reactome highlighting neuronal signaling and cholesterol biogenesis and KEGG representing specific neurotransmitter and endocrine signaling processes within the downward DEG’s. Interestingly, KEGG pathway enrichment identifies GABAergic synapse down-regulation, consistent with recent findings suggesting that loss of GABAergic neurons occurs earlier in the disease sequelae(48). Conversely, the up-regulated DEGs enrich cell cycle and DNA replication and repair GO biological processes, while KEGG pathway enrichment highlights complement and coagulation cascades, pointing towards innate immune processes involved in synaptophagy (Figure 1D, right side, second and third rows).

The TREAT-AD biological domains map each dataset independently, both murine and human, into the 19 discrete endophenotypic areas (Figure 2). The Kat5 model (upper left) upregulates cell cycle, immune response, and structural stabilization (a domain reflecting cell-cell junctions, connections to cytoskeletal elements and ECM factors(33)), while downregulating synapse, endolysosome, myelination and mitochondrial metabolism (Figure 2). The murine 5XFAD model, also upregulates immune response and downregulates synapse (upper right, Figure 2). The human datasets representing three distinct AD subtypes: LOAD, EOAD (non-genetic), and FAD (PSEN1 mutation). Each subtype demonstrates upregulated immune response, cell cycle, DNA repair, and structural stabilization, concomitantly with downregulated synapse, mitochondrial metabolism and endolysosome, all shared with the Kat5 AD biological domain profile. A set of biological domain specific GO-terms are enriched across all datasets, including positive enrichment of complement activation and negative enrichment of synaptic vesicle recycling and postsynaptic density.

**Figure 2.**
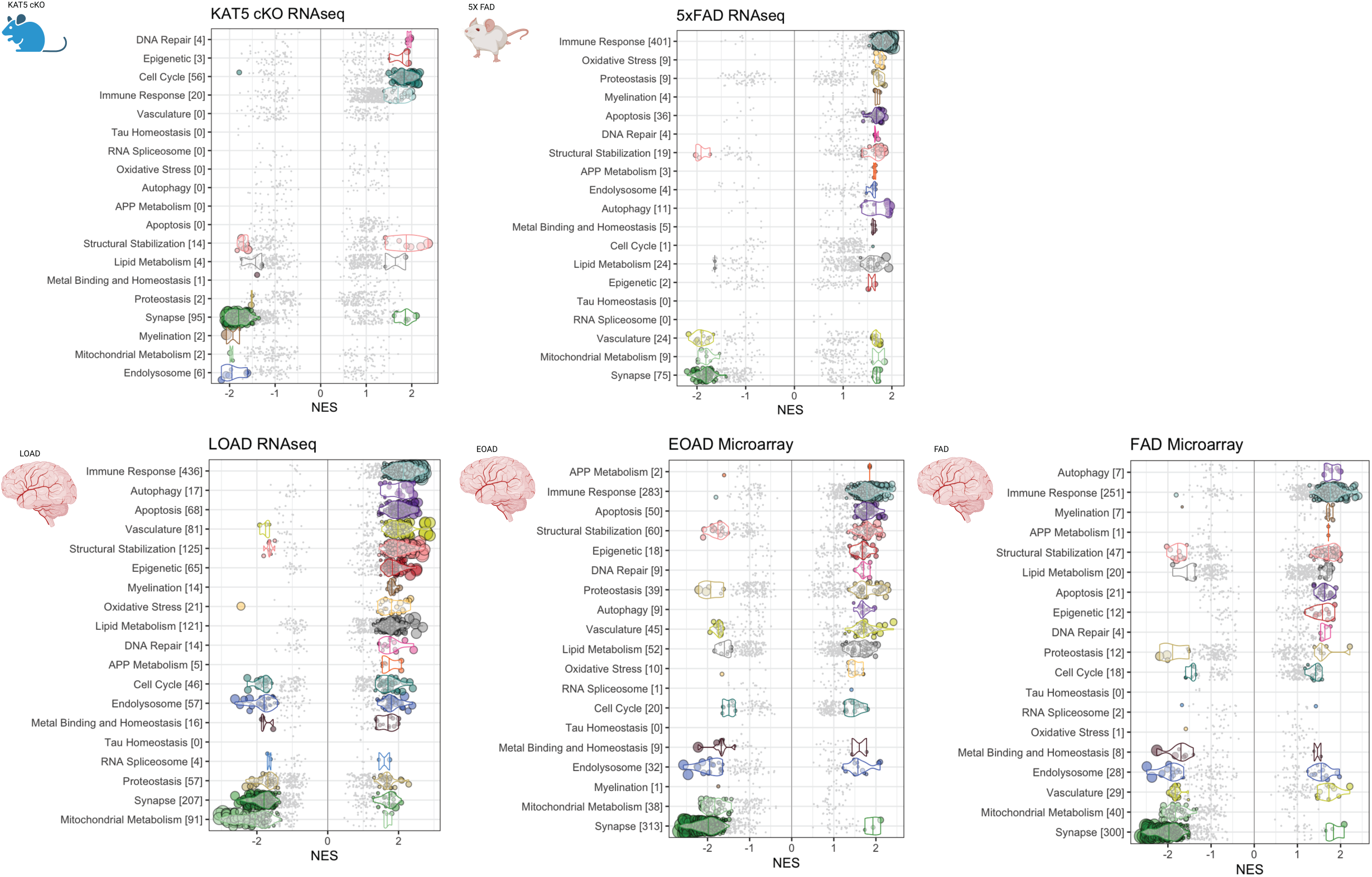
Biological Domain Comparisons of All AD Datasets. All datasets including KAT5, the 5X FAD murine model, human LOAD transcriptome, EOAD and FAD human datasets are broadcast onto the TREAT-AD biological domains based upon GSEA enrichment of GO terms within the DEGs of each transcriptome. The murine models are on the top, denoted by mouse symbols, while the 3 human datasets are on the bottom: LOAD (left), EOAD (middle), and FAD (right). The negative normalized enrichment score (NES) denotes downregulated GO terms, while positive NES means the leading-edge genes associated with that term are upregulated.

The methodology employed for Kat5 and AD dataset comparisons is shown diagrammatically in Figure 3A, with the pairwise analysis shown on the left, and the methodology employed shown to the right. The Pearson correlation (r) associated with each binary model comparison demonstrates a relatively weak correlation overall (ranging between 0.11 for LOAD and 0.35 for 5XFAD), but highly statistically significant (Figure 3B), suggesting overlap within a portion of the DEGs. To hone in on the biological areas of elevated correlation within the Kat5 and AD comparisons, we examine the overlap within the AD biological domains.

**Figure 3.**
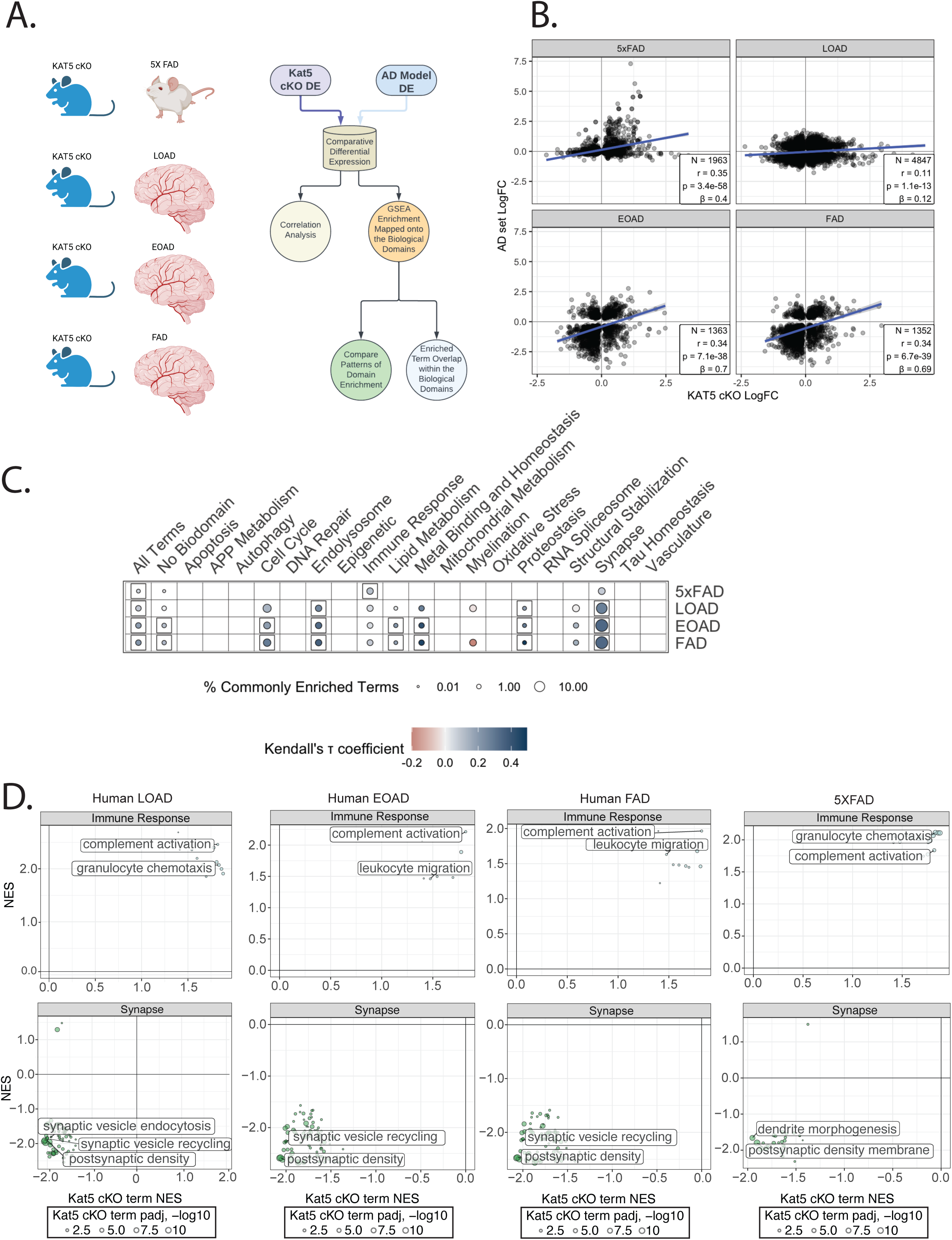
KAT5 correlations with AD datasets. To test the similarity of the differential expression pattern observed within the KAT5 cKO to extant AD datasets, we performed a set of comparisons to murine and human AD subtypes, with the diagrammatic process depicted in (A), the Pearson correlations shown in (B) and the correlation between each dataset with KAT5 DEGs mapped onto the TREAT-AD biological domains (C). The binary correlations between KAT5 and each AD subtype or model are provide for the most consistently up and down-regulated biological domains, immune response and synapse respectively (D).

The AD biological domains are a TREAT-AD resource developed to subdivide gene sets into specific biological endophenotype-linked processes(33). Consequently, comparing the correlation within specific AD biological domains should show the AD linked biology with highest correlation between datasets. Across all examined murine and human datasets, synapse and immune response are the most consistently and positively correlated biological domains (Figure 3C). There is significant correlation between Kat5 and all human datasets within the endolysosome and proteostasis domains.

The Kat5 cKO comparison to the LOAD DEG dataset is critical to understanding if Kat5 plays a role in AD pathogenesis, and if so, within which biological processes or pathways. To examine this issue, we intersect the downregulated genes from LOAD and Kat5 cKO and use the 845 shared downregulated geneset to seed GO and pathway enrichment analyses (Figure S1, top, Venn diagram). The GO enrichment of the common downward DEGs highlights synaptic processes, including presynaptic and postsynaptic processes and pathways (Figure S1, bottom panels). There is notable enrichment of GABAergic signaling (Figure S1: KEGG, WikiPathways, GO cellular component), suggesting a shared impairment in inhibitory synapses within LOAD and the Kat5 cKO model. Neurodevelopmental processes are represented within the enrichment analyses, such as axonal projection or neurodevelopmental process (Figure S1: GO BP CC, Reactome), which is consistent with the kappa-network enrichment of the KAT5 model (Figure 1). Metabolic pathway downregulation of glycolysis and lipid metabolism (Figure S1: WikiPathways, MSIGDB) is common to LOAD and Kat5 cKO, further supporting metabolic impairment as a common biological process disrupted across datasets. Finally, LOAD and Kat5 cKO data share signatures with the GTEX genes downregulated in brain between young subjects (20–29) and older individuals (60–79), arguing for LOAD and Kat5 cKO convergence on aging brain genes (Figure S1, bottom right).

The 599 upregulated DEGs intersecting LOAD and Kat5 cKO are employed to seed reciprocal enrichment analysis of these two datasets (Figure S2, Venn diagram). The GSEA of the common DEGs demonstrate GO term and pathway enrichment of immune function, DNA replication and repair. The common immune genes are involved in proinflammatory cytokines, such as TNF and members of the interleukin family, as well as phagocytic and complement pathways (Figure S2: KEGG, WikiPathways). GTEX signatures show brain aging genes are upregulated (Figure S2), converse to the downregulation of GTEX brain aging signatures in downregulated DEGs. We examined the gene overlap within the discordantly regulated gene sets to assess if there are processes that were markedly opposed between LOAD and the Kat5 cKO, which pointed toward transcriptional or chromatin regulation within Kat5 Down-LOAD Up geneset and DNA repair or cell cycle processes within the Kat5 Up-LOAD Down gene sets (Figure S3).

We performed supplementary analysis of the downregulated gene intersection between LOAD and Kat5 DEGs in SynGO, a portion of the GO focused upon synaptic structure and process. This identified pre- and postsynaptic processes (Figure S4), with similar degrees of representation, showing dendritic and axonal processes in the overlapping Kat5 and LOAD biology. Consistent with the biological domain analysis, the metabolic profile of the overlapping downregulated genes implicates GABA, as well as NAD and NADH (Figure S4, Human Metabolome Database). This metabolic data supports Kat5 and LOAD dysregulation of GABA-related signaling and the TCA cycle, which fuels the electron transport chain (ETC), further supporting the decrement in synaptic and mitochondrial function as common biological impairments.

Examining specific gene expression changes in Kat5 cKO, we observe downregulation of neuronal glucose transport (Glut3/Slc2a3) and glycolytic metabolism of glucose into pyruvate (Figure S5A). Mitochondrial genes involved in pyruvate import into mitochondria and TCA cycle function are also downregulated (Figure S5B), along with key electron transport chain genes (Figure S5C). These observations further demonstrate mitochondrial impairments in the Kat5 cKO mouse model. We explored whether upregulation of immune response in Kat5 cKO might impinge upon microglial state regulatory genes. These analyses show small changes within homeostatic marker genes in microglia, particularly P2ry12 (Figure S6A), and upregulation of many activation markers, including Syk, TyroBP, Axl, and Trem2 (Figure S6B). The Kat5 gene deletion occurs only in neurons, which argues for a non-cell-autonomous effect of neuronal deficits on microglial activation.

To assess the role of KAT5 in the central nervous system, outside of a disease context, we utilized the CRISPRbrain resource(49). CRISPRbrain employs human induced pluripotent stem cells (hiPSCs) differentiated into neurons, astrocytes and microglia to interrogate functional genomics within assays of survival, metabolic tone, ROS production, phagocytosis, and immune activation, among others. KAT5 inhibition consistently decreased neuronal survival and was the most significant epigenetic regulator for neuronal survival (Figure 4A, purple=impaired survival; green=promoted survival). Interestingly, within histone acetylases and deacetylases, gene deletion of either class has a predominantly negative impact on neuronal survival, suggesting a complex role for epigenetic regulation in neuronal survival outcomes. Loss of KAT5 or CREBBP, both major brain histone acetylases, diminish neuronal survival (Figure 4A).

**Figure 4.**
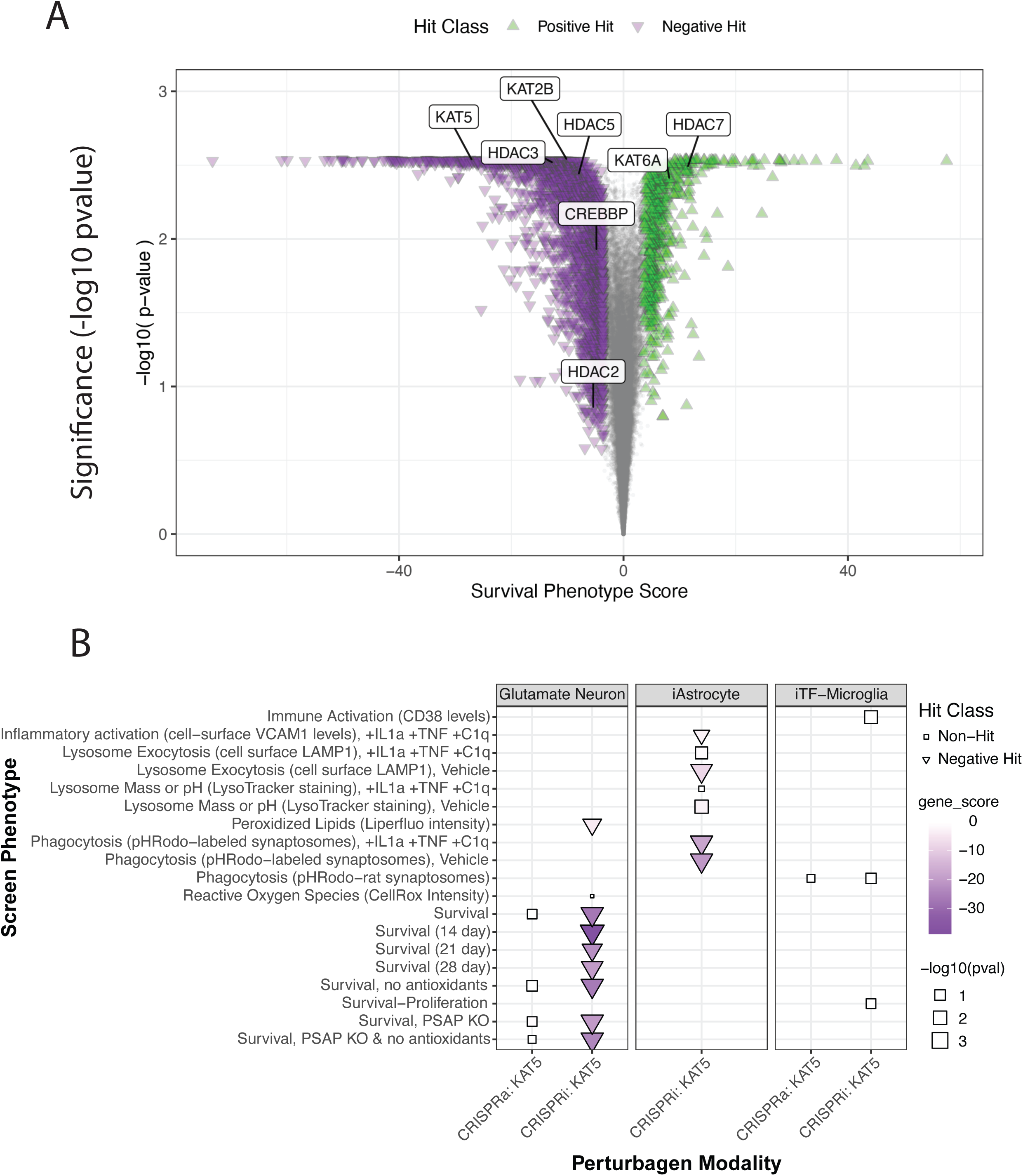
KAT5 and epigenetic regulatory factors in involved in neuronal survival in CRISPRbrain studies. CRISPRbrain performs functional genomics studies in iPSC-derived brain lineages in association with focused assays examining traits of relevance for human health such as neuronal survival. The CRISPRbrain genetic models employ both CRISPRi and CRISPRa approaches to assess the impact of up or downregulation of a specific gene upon cellular physiology. (A) The neuronal survival assay is demonstrated by volcano plot using survival assay phenotype score along the abscissa and the probability along the ordinate axes. KAT5 is the most significantly ranked epigenetic regulator, the deletion of which having negative impacts upon the neuronal survival score. Important to note, other epigenetic regulators with shared and opposing function are also linked with neuronal survival. (B) The collective set of simple assays performed with the KAT5 CRISPRi model in different cellular lineages are plotted, with intensity of color denoting the gene score associated with the assay, the size of the icon denoting statistical significance, the triangle shape associated with a negative hit, while a square shape denoting a non-hit. All neuronal survival assays performed across different timeframes show KAT5 deletion negatively and significantly associated with survival. Phagocytotic astrocytes are also negatively associated with KAT5 function. There are no significant microglial assay results for KAT5.

Panning across all CRISPRbrain assays performed in all three cell types, we observe a uniformly consistent decrement of neuronal survival in KAT5 knockdown neurons (Figure 4B, left box, CRISPRi), with little effect of KAT5 activation (Figure 4B, CRISPRa). In astrocytes, however, inhibition of KAT5 produced notable deficits in lysosome exocytosis, inflammatory activation in response to cytokine (TNF/IL-1a) and complement (C1q) treatment, as well as significant impairment in synaptic phagocytosis (Figure 4, middle box). These observations support a role of KAT5 in multiple cell-type specific processes involved in the AD pathogenic sequalae.

To explore genetic support for the transcriptomic profiles of Kat5 cKO, genetic variant trait-linkages was performed with the up- and downregulated genes using data from the 2023 GWAS catalogue, which contains over 400,000 SNP-trait associations across greater than 5000 associated traits(41). The analysis is performed in a stepped manner, in which we pan across the downregulated gene set based on the degree of differential expression. The goal is to balance potential issues of sensitivity of enrichment, that may be diluted within the lower level differentially expressed genes, versus truncating implicated processes by setting too stringent a standard based on high fold-enrichment. Within the downregulated gene set, the primary enrichment employs the 2792 DE genes that are 1.2-fold downregulated (most permissive) in the Kat5 cKO model. When we perform the trait enrichment, the top terms imply linkage with anatomical development, cognitive and psychiatric health (biopolar disorder, highest math score), as well as direct trait-linkage with AD (‘Alzheimer’s Disease (Cognitive Decline)’, ‘Psychosis and Alzheimer’s Disease’) (Figure 5A). Increasing the stringency to 1.5-fold DEGs (which number 949), we observe the top two enriched term are ‘Psychosis and Alzheimer’s Disease’ and ‘Lateral Ventricle Volume’, a known physiological biomarker of limbic and temporal lobe shrinkage associated with neurodegeneration (Figure 5B). At 2-fold DEG, the highest stringency tested, we have 168 genes trait-linked with ‘diabetes medication use’, ‘central cerebrospinal fluid amyloid Aβ42 to Aβ40 ratio’, and ‘lateral ventricle horn volume’, all possessing directly or indirect associations AD pathological progression(1, 50). A similar analysis was performed with the up-regulated genes with no compelling findings (Figure S8). Consequently, the downregulated genes within the early developmental stage, neuron-specific Kat5 induced forebrain mouse knockout are genetically associated with human traits directly connected to AD, making a strong case for a direct role for KAT5 in AD pathological progression.

**Figure 5.**
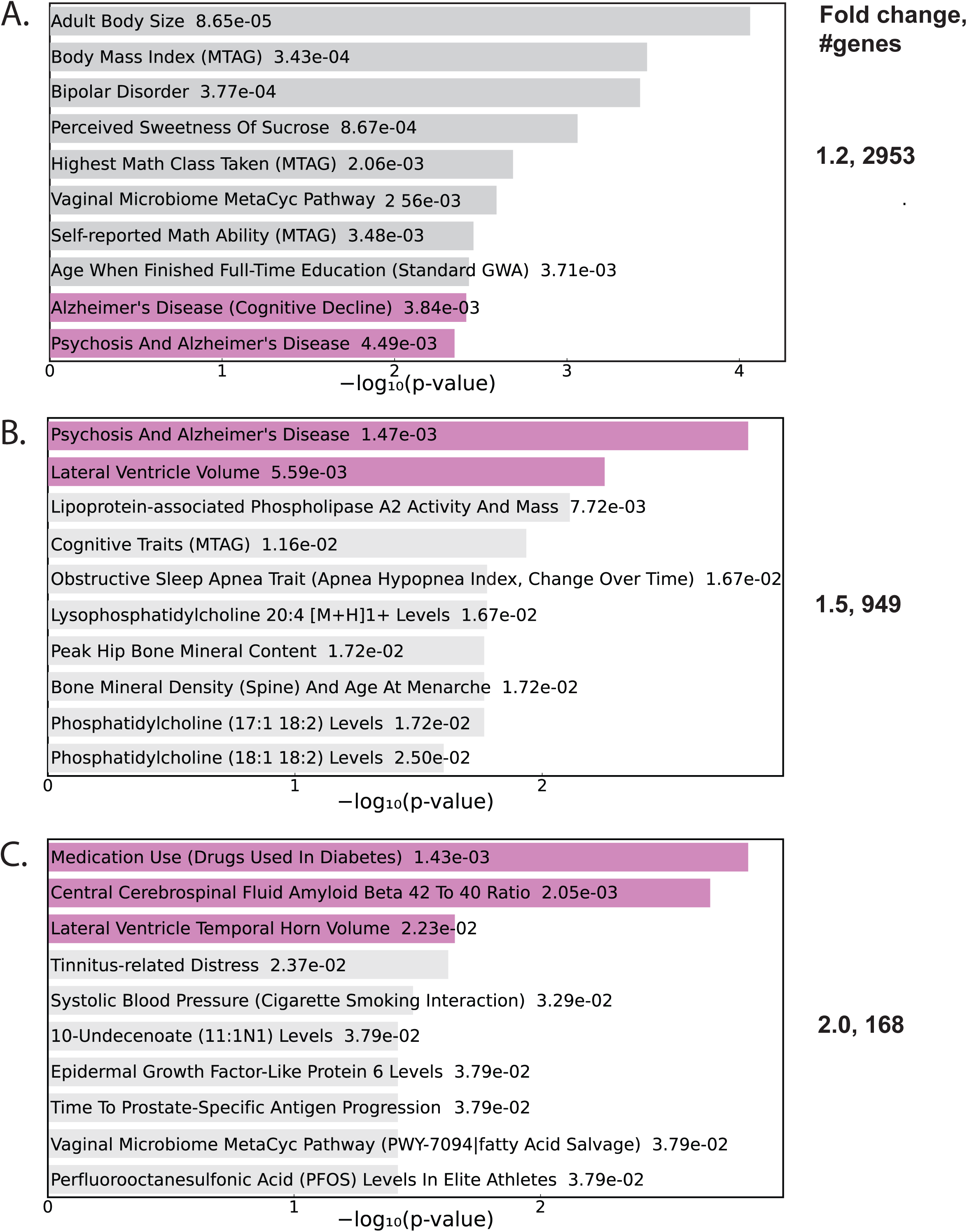
Genetic Trait Mapping of KAT5 Downregulated Genes. The GWAS 2023 Catalogue was trait mapped using three different stringencies of the KAT5 downregulated genes: mild (1.2 fold), moderate (1.5 fold) and strong (2 fold). The more permissive DEG filtering result in different size gene sets ranging from 2953 (mild), 959 (moderate) and 169 (strong). The KAT5 DEGs were matched to the GWAS catalogue genes and the traits to which variants demonstrated linkage. In all three stringencies of filtration, the KAT5 cKO attenuated DEGs map onto AD, or AD-related traits. In the mild condition, AD (Cognitive Decline) and Psychosis and Alzheimer’s Disease are observed. In the middle analysis, the top two enriched traits are Psychosis and Alzheimer’s Disease and Lateral Ventricle Volume (a physiological biomarker of AD). In the most stringent analysis shown in the bottom panel, we observe enriched trait linkage with Medication Usage (Diabetes), Cerebral Spinal Fluid Amyloid 42 to 40 Ratio, and Lateral Ventricle and Temporal Horn Volume.

The fundamental hypothesis of this work is that a decrement in gamma-secretase function associated with AD (FAD or LOAD) may contribute to differential subcellular localization of KAT5 through membrane sequestration due to its association with APP and FE65/APBB1. To directly test this idea, we examined iPSC-derived neurons in wild-type and AD lineages. In wild-type human neurons, we observe a relatively diffuse state of KAT5 localization throughout the cell (Figure 6A). The DAPI staining (blue) demonstrates the nuclear boundaries, which KAT5 enters within the WT neurons. The examination of KAT5 in heterozygous APP_swe_ (Figure 6B, neurons isogenic to the WT with CRISPR gene editing insertions of the APP Swedish heterozygous mutations(51)) demonstrate cytosolic restriction. Similarly, KAT5 is restricted from the nucleus in the APP_dup_ (Figure 6C, neurons carrying a duplicate APP allele) human neurons. The examination of the PSEN2 FAD heterozygous mutant neurons also demonstrate a predominantly cytoplasmic sequestration (Figure 6D). Concordant with the shift in localization being due to a diminution of gamma-secretase activity in the mutant neurons, when DAPT is applied to the WT neurons, KAT5 is restricted from the nuclear compartment (Figure 6E), suggesting a gamma-secretase mediated event is required for nuclear localization of KAT5.

**Figure 6.**
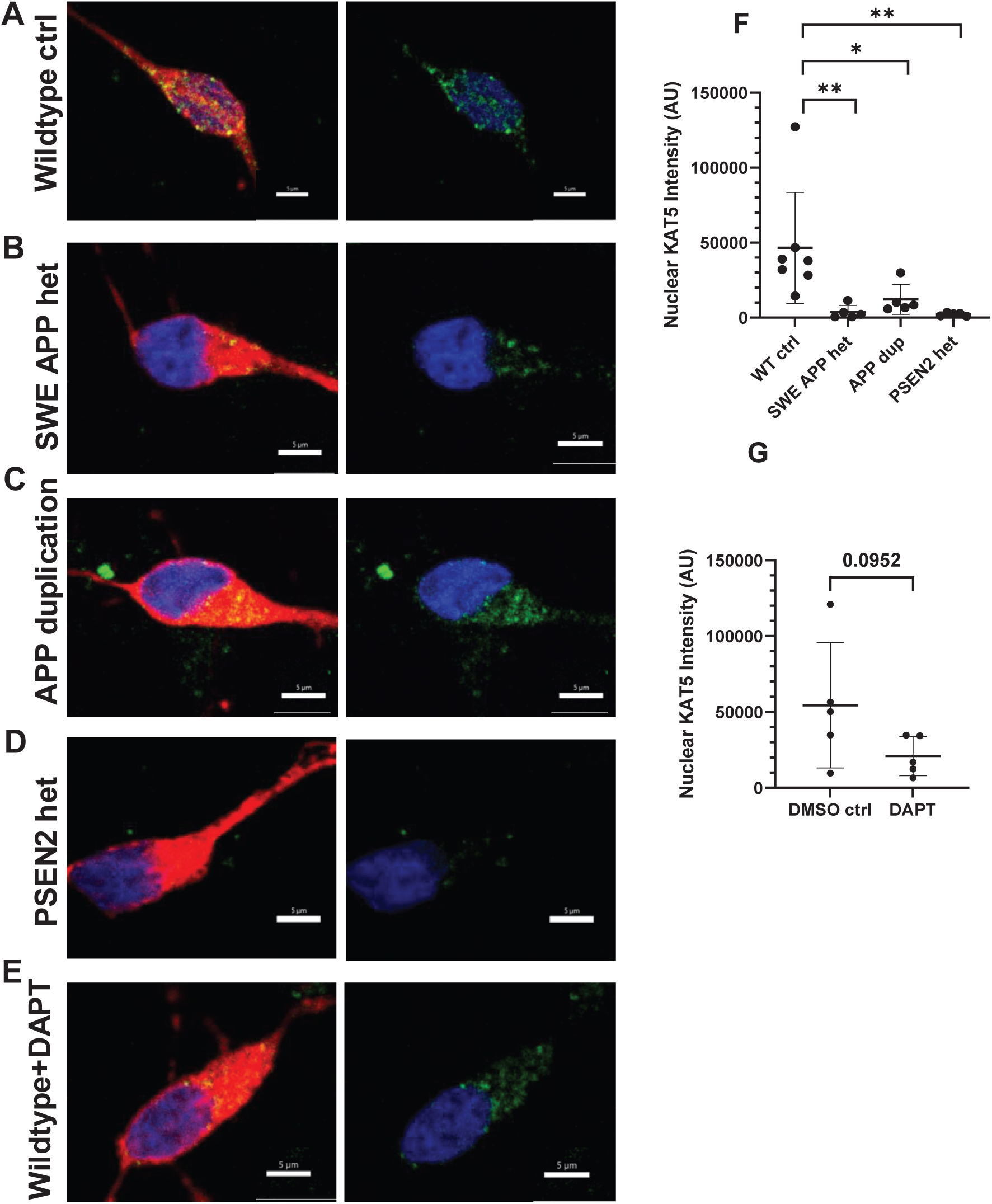
Subcellular localization of KAT5 in human iPSC-derived WT and FAD neurons. Induced-pluripotent stem cells (iPSCs) from a WT CViia2 strain, APP Swedish (APPswed) and APP duplication mutants (APPdup, a model of trisomy 21) were differentiated into glutamatergic neurons and stained for KAT5 (green), DAPI (blue) to show nuclear localization and MAP2 (red) to show cytosolic cytoskeletal structure. The wild-type cells demonstrate ubiquitous localization (A) with fluorescent overlap with the DAPI nuclear signal. All explored FAD mutant lines (B and C) and APP duplication lines (D) show specific nuclear exclusion of KAT5. The WT CViia2 neurons treated with gamma-secretase inhibitor DAPT demonstrate nuclear exclusion of KAT5 (E). The statistical significance was analyzed as described in methods section across cells per culture group, with the average for each cell-type shown as a single dot in the plots on the right (F and G). The WT cells are statistically significantly different in nuclear signal intensity than either the APPSWE heterozygous cells (p<0.01), the APP duplication (p<0.05), and PSEN2 N141I heterozygous mutants (p<0.01). The gamma-secretase inhibitor treated cells approached statistical significance (p<0.09).

Based upon the MAP2 staining (red), all neurons appear healthy and viable. The images were quantitated as described in the methods section and show significant decreases in nuclear KAT5 staining intensity (Figure 6F). The quantification of the DAPT treated cells show a comparable decrement in nuclear intensity that is significant at p<0.1 level (Figure 6G). Of interest, the staining in the APP_swe_ and APP_dup_ lines appears largely punctate, likely due to vesicular trafficking. A previous study demonstrated that FAD mutations in APP and PSEN1 induce endolysosomal enlargement and aberrant trafficking of APP(52). Our data is consistent, potentially noting that KAT5 remains associated during the endolysomal pathological progression. These data directly support the central hypothesis of this work.

To further explore the plausibility of KAT5 driving elements of AD pathology, we examined the single cell expression patterns of AD brains characterized in the recent SEA-AD study(48). Consistent with reports of FAD mutations inducing an enzymatic stalling of the gamma-secretase complex with the APP substrate(21), we observe down-regulation of some components of the heterotetrameric gamma-secretase complex across AD progression, modeled within the SEA-AD study by a neuropathologically derived continuous pseudoprogression score (CPS) (Figure 7). We specifically observe decrements in PSEN2 (Figure 7B) and APH1B (Figure 7D) within both the inhibitory and excitatory neuronal subtypes (left) and supertypes (right). Additionally, there is a decrement in NCSTN (Figure 7E) in the later stage of AD progression, particularly within the inhibitory neuronal populations. Potentially compounding KAT5 dysregulation, we also observe that in GABAergic and glutamatergic neurons, there is a decrement in KAT5 expression across AD pseudo-progression (Figure S7). The decrease in KAT5 expression appears in the early and late stages of disease, in neuronal subtypes (Figure S7A) and supertypes (Figure S7B). This may trigger early epigenetic changes preceding the severely detrimental late-stage progression in disease sequalae identified within the SEA-AD study. Given the upregulation of APP in AD human post-mortem brain(53), the downregulation of KAT5 could result in an elevated proportion of KAT5 localized to the membrane. The decrement in KAT5 expression is present in most neuronal subtypes, including the somatostatin inhibitory neurons, that the study identified as one of the earliest neuronal lineages to be impacted by AD pathogenesis(48).

**Figure 7.**
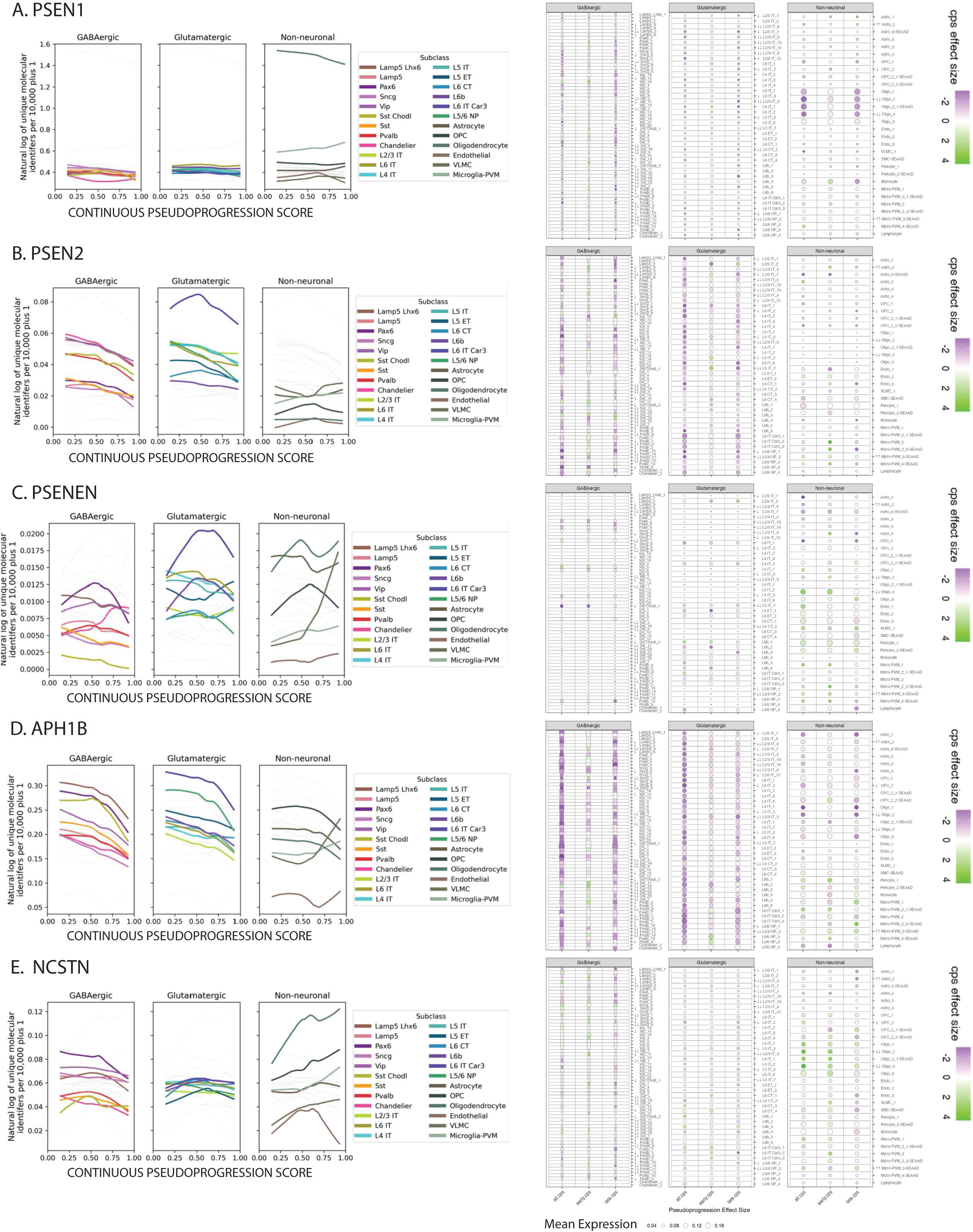
Gamma-Secretase Components Dysregulated in Neurons in SEA-AD LOAD Study. The Seattle Alzheimer’s Disease (SEA-AD) study performed expression analysis across AD progression as modeled by the continuous pseudo-progression score (CPS) within all identified cellular subtypes within the human brain taxonomy. The gamma-secretase components are shown at two levels of cellular resolution with column on the left depicting subtype (more general) cell types and the column on data on the right hand side showing all available cellular subtypes. Both views divide the cellular lineages into inhibitory GABAergic neurons, excitatory glutamatergic neurons, and non-neuronal lineages (glial and endothelial subtypes). The representations on the left quantitate expression across disease based upon normalized expression levels and on right the expression is heat-mapped based on CPS effect size with down-regulation represented in purple and upregulation represented in green and the size of the circle representing mean expression level. The homologues of presenilin PSEN1(A) and PSEN2(B) levels across disease are shown in the top two rows. The remains three components of the heterotetrameric gamma-secretase complex are shown below (C) PSENEN, (D) APH1B, and (E) NCSTN. At the low (left) cellular specificity and high cellular specificity (right) representations, there are strong downward regulation of PSEN2 and APH1B within both inhibitory and excitatory neurons.

The Kat5 cKO overlap in molecular profile with FTD patient transcriptomic data from the RiMOD consortia (see methods) was explored to assess the AD specificity of the similarity. The overlap between Kat5 cKO and the FTD patients was considerably smaller than with AD (Supplementary Figure 9), with less than 10% overlapping genes in the up or down-regulated gene set. Not surprisingly, there are some similarities in biological enrichment, that likely owe to common molecular pathways involved in neurodegenerative events, also observed in recent explorations of the molecular profiles across neurodegenerative diseases(54).

## 4. Discussion

The core question investigated within this work is whether KAT5 mediated epigenetic regulation could be involved in normal neurological homeostasis, the disruption of which may dysregulate CNS processes and catalyze the neurodegenerative cascade observed within AD. KAT5 is known to be critical in development, as the Kat5 knockout mouse is embryonic lethal in the early blastocyst stage(55) and CNS homozygous ablation of KAT5 leads to malformation of cortical tissue and microencephaly(56), suggesting a prominent role for epigenetic regulation in early central nervous system development. Identified hypomorphic missense variants in KAT5 are linked with cerebellar encephalopathy and decreases in cognitive function within the variant carriers, further supporting the linkage between KAT5 and neurological development(57). The purported mechanism underlying the epigenetic hypothesis presented here is that KAT5 association with APP and FE65/APBB1 at the synaptic membrane gates its nuclear signaling through cytoplasmic sequestration.

To address this hypothesis, we have employed a re-analysis of the CNS specific inducible knockout of the Kat5 mouse gene(31), aligning it to the AD biological domains developed within the TREAT-AD consortia, and comparing it to multiple other AD transcriptomic datasets derived from patient brain banks and a relevant AD mouse model (Figure 3A). The primary impetus for this investigation comes from a longstanding observation that many of the familial Alzheimer’s disease (FAD) causing mutations diminish the efficacy of gamma-secretase mediated APP proteolysis(14, 15, 58, 59). One plausible mechanism to explain the deficit in enzyme function in the face of elevated amyloid peptide, revolves around the processivity of gamma-secretase(15, 19). In this model, decreases in proteolysis correspond to longer and more pathogenic amyloid peptides(19). FAD mutations in APP and PSEN1 result in a stalling of the enzyme-substrate complex at the transition state intermediate, preventing completion of the proteolytic event(21), which dramatically reduces the liberation of the APP intracellular domain (AICD), and may trap KAT5 at the synaptic membrane.

The primary prediction of this model is that KAT5 membrane sequestration may alter nuclear chromatin regulation impairing essential neuronal homeostasis and contribute to AD pathogenesis. In our characterization of the Kat5 conditional knockdout (cKO), we observe a profound synaptic phenotype, as systemic downregulation of synaptic gene expression is the core transcriptomic signature (Figure 1, Figure 3). Intriguingly, we observe downregulation in both excitatory and inhibitory neuronal pathways in the Kat5 cKO, the LOAD transcriptomic dataset(33), and within downregulated genes common to both (Figure 1, Figure S1). We also observe that disruption of Kat5 results in decrements in synaptic developmental signatures (Figure 1A, B, D), in support of the above hypothesis. The upregulated DEGs in the Kat5 mouse implicate cell cycle, DNA repair and immune response (Figure 1A, C, D, Figure 2), signatures that are also present in LOAD (Figure S2).

The direct comparison of the Kat5 cKO mouse model transcriptomic signature to LOAD, early onset AD [not associated with FAD mutations] (EOAD), FAD, and the 5X FAD mouse (Figure 3A) show overlapping patterns of biological domain enrichment in synapse, immune response, cell cycle and others (Figure 2, 3). The correlation between Kat5 cKO DEGs and the comparative AD datasets is weak but highly significant (Figure 3B), due to the elevated correlation observed in specific overlapping biological domains (Figure 3C), most consistently synapse and immune response, with overlap between Kat5 cKO and the human datasets observed in cell cycle and endolysosome domains as well (Figure 3C). Intriguingly, we observe upregulation of complement cascades and immune migration pathways activated across all models (Figure S3, Figure 3D) which is consistent with the shifts in the Kat5 cKO from homeostatic to activated disease associated microglial markers (Figure S6). Surprisingly, there was little overlap in mitochondrial function, the biological domain in which we observe the greatest reduction in LOAD(33), but we do see focal downregulation of a multitude of metabolic pathways involved in glucose transport, glycolysis, TCA cycle and ETC (Figure S5). These observations support a role for KAT5 in regulation of metabolic genes. Interestingly, KAT5 is part of the NuA4 complex, that associates with the transcription co-activator JAZF1, which targets KAT5 to the promoters of metabolic genes, enhancing subsequent transcription(60, 61). Concordantly, JAZF1 is independently identified as an AD genetic risk factor emerging from recent GWAS studies(62). KAT5 epigenetic regulation of genes involved in AD pathophysiology is consistent with the observation that the Kat5 cKO downregulated DEGs demonstrate strongly enriched trait-variant linkage with cognition and AD pathology across tiers of differential expression (Figure 5); no such relationship is observed with the upregulated genes (Figure S8).

The role of KAT5 in CNS physiology is likely strongest in neurons, as KAT5 repression through CRISPRi (49) robustly inhibits neuronal survival within CRISPRbrain studies (Figure 4A, 4B left panel). Surprisingly, we did not observe any microglial associated impacts of KAT5 inhibition within the assays performed, which supports the non-cell autonomous microglial activation by KAT5 deletion in neurons in the Kat5 cKO mouse. In astrocytes, however, KAT5 repression elicits a decrement in phagocytosis of synapses, supporting cell lineage specific effects (Figure 4B), and suggest further investigation of KAT5’s role in astrocytes within glial-targeted mouse models would be warranted in the future.

The gamma-secretase dependence of KAT5 nuclear signaling was explored in subcellular localization studies in human wild-type and AD iPSC-derived human neurons. We compared KAT5 localization in human iPSC-derived neurons from wild-type (WT) and those carrying the APP Swedish, an APP duplication, and PSEN2 N141I FAD mutations. The WT cells demonstrate a ubiquitous distribution of KAT5 throughout the cell, while neurons treated with DAPT (a gamma-secretase inhibitor) (Figure 6E) or cells carrying FAD mutations show punctate cytosolic sequestration (Figure 6). These data are consistent with previous reports of nuclear exclusion in human hippocampal neurons in post-mortem AD brains(30). Further, previous studies of large panels of FAD iPSC-derived neurons, show consistent endolysosomal enlargement and increases in APP CTFs trafficked to the endolysosome(43, 52, 63), similar to the punctate pattern observed with KAT5 in this study (Figure 6). FAD mutations may trigger endolysosomal trafficking of APP in which KAT5 remains membrane tethered. The potential relevance to LOAD is noted within the reanalysis of the SEA-AD study, in which core components of the heterotetrameric gamma-secretase complex (especially PSEN2 and APH1B, Figure 7) are down-regulated in LOAD along with KAT5 itself (Figure S7). These data support the idea that increases in APP and relative decreases in gamma-secretase mediated liberation of KAT5 may be involved in both FAD and LOAD. This conjecture is further supported by the enrichment of GTEX brain aging signatures within the Kat5 cKO intersection with LOAD (Figure S1 (common downregulated), Figure S2 (common upregulated). Additionally, we examined the AD specificity of the molecular signatures comparing the Kat5 KO transcriptome to FTD patients in the RiMod consortia, which demonstrates in a weak overlap in altered gene expression (Supp Figure 9); this is consistent with the down-regulated genes showing genetic linkage with AD specific processes (Figure 6).

There are limitations in this analysis, both in terms of the biological models and the analytic methodology. One extreme limitation is the deployment of transcriptomic sequencing in a very early developmental stage mouse (3 weeks postnatal), which precedes any observed pathology, and may minimize the impact on the emergent molecular signatures. Future studies will examine a broader range of developmental stages. Alternative interpretations are also possible, as KAT5 may be involved in core neurodevelopmental or neuronal homeostatic processes that are independent of AD, or not specific to the disease state. This possibility cannot be fully ruled out, but the similarity of AD subtypes and murine model transcriptomic signatures support mechanistic conservation, pointing towards direct involvement of KAT5 in the AD neurodegenerative sequalae.

### 4.1 Conclusion

Collectively, the observations made within this analysis support a model of AD pathogenesis in which APP associates with APBB1/FE65 and KAT5 at the synaptic membrane of neurons, and following gamma-secretase mediated APP cleavage, liberate KAT5 to migrate to the nucleus and activate homeostatic renewal of synaptic and mitochondrial genes (Figure 8A, C). This model supposes that APP/KAT5 signaling is utilized in development to facilitate synaptic wiring in the brain and shifts to homeostatic regulation in the mature organism, which is supported by the strong neurodevelopmental enrichment patterns within LOAD and the Kat5 mouse model. The Kat5 cKO stimulates massive gliosis of both astrocytes and microglia by nine months of age(31), and even at two-weeks post induction of the Kat5 knockout, we observe upregulation of numerous immune-related targets (Figure 2, Figure S2, Figure S6), supporting the idea that KAT5 deficiency could promote a transition between homeostatic and activated microglia (Figure 8B). KAT5 cytoplasmic sequestration is observed in human AD neurons (Figure 6) and has been observed within LOAD hippocampal tissue(30), and alteration of the KAT5/HDAC2 levels are associated with increased cognitive performance(29, 30), which is consistent with the Kat5 subcellular localization to the nucleus being critical to learning in drosophila(64). Consequently, the plausible role of APP/KAT5 signaling dysregulation contributing to AD pathogenesis is supported by common transcriptomic patterns in the molecular signatures observed in biological domains and pathway analysis but will require future investigation to determine the validity of this mechanism in promoting AD pathogenic progression.

**Figure 8.**
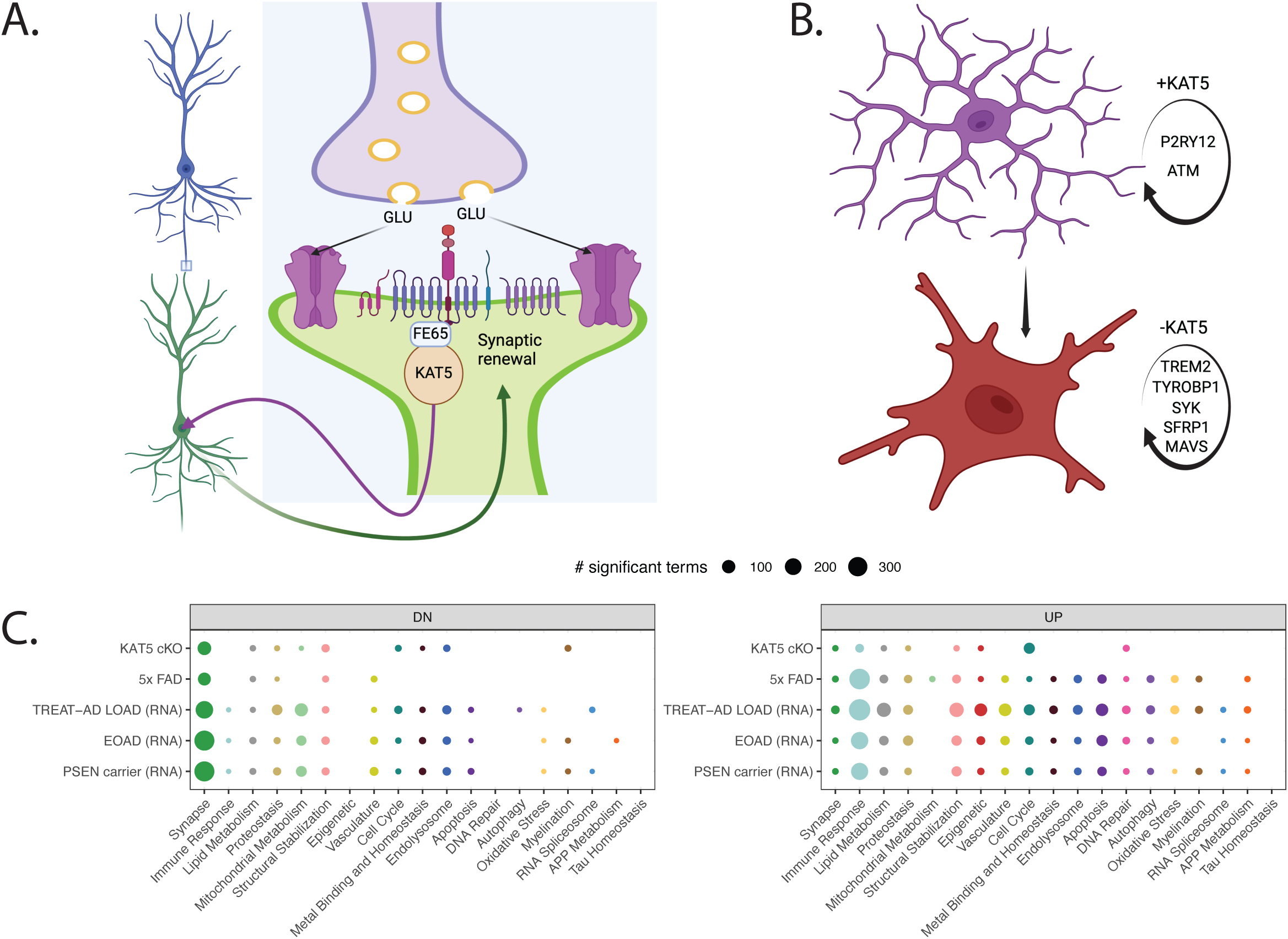
Model of Hypothetical KAT5 Biological Function in Neurons and Microglia. (A) The enrichment data points to a developmental program in which KAT5 promotes synaptic gene expression potentially fostering synaptic development. KAT5 is known to associate with APP via the APBB1/FE65 binding protein and following gamma-secretase mediated cleavage may travel back to the nucleus to stimulate a renewal wave of synaptic gene expression. (B) Numerous genes associated with microglial activation state are regulated by KAT5 expression, either directly or indirectly, suggesting that KAT5 may be involved in maintaining microglia in a homeostatic state. Consequently, a lack of KAT5 nuclear signaling would be predicted to decrease synaptic gene expression and increase immune response genes associated with reactive gliosis. (C) Comparing all the models explored in this study across all 19 biological domains for both downregulated and upregulated gene sets, we see a consistent decrease in synaptic signal across all models, including KAT5, as well as increases in immune response, consistent with the hypothesis suggested in (A) and (B). (A) and (B) were generated using BioRender software.

## 5. Data Availability/Use

The LOAD dataset employed in this work is derived from TREAT-AD harmonization and resource development work, and is available on the AD Knowledge Portal (https://adknowledgeportal.synapse.org; synapse ID: syn21983020). The EOAD and FAD datasets are available on GEO at accession GSE39420. The Kat5 cKO dataset is available in association with the original publication(31).

## Abbreviations

(AMP-AD): Accelerating Medical Partnership for AD
TREAT-AD): Target Enablement to Accelerate Therapy Development in AD
(MODEL-AD): Model Organism Development and Evaluation for Late-onset AD
(GSEA): Gene Set Enrichment Analysis
(GO): Gene Ontology
(CADRO): Common Alzheimer’s and Related Dementias Research Ontology
(TRS): Target Risk Score

## References

1. Knopman DS, Amieva H, Petersen RC, Chételat G, Holtzman DM, Hyman BT, et al. Alzheimer disease. Nat Rev Dis Primers. 2021;7(1):33.

2. Wolfe MS. Unraveling the complexity of γ-secretase. Semin Cell Dev Biol. 2020.

3. Szaruga M, Munteanu B, Lismont S, Veugelen S, Horré K, Mercken M, et al. Alzheimer’s-Causing Mutations Shift Aβ Length by Destabilizing γ-Secretase-Aβn Interactions. Cell. 2017;170(3):443–56.e14.

4. von Koch CS, Zheng H, Chen H, Trumbauer M, Thinakaran G, van der Ploeg LH, et al. Generation of APLP2 KO mice and early postnatal lethality in APLP2/APP double KO mice. Neurobiol Aging. 1997;18(6):661–9.

5. Herms J, Anliker B, Heber S, Ring S, Fuhrmann M, Kretzschmar H, et al. Cortical dysplasia resembling human type 2 lissencephaly in mice lacking all three APP family members. EMBO J. 2004;23(20):4106–15.

6. Korte M, Herrmann U, Zhang X, Draguhn A. The role of APP and APLP for synaptic transmission, plasticity, and network function: lessons from genetic mouse models. Exp Brain Res. 2012;217(3-4):435–40.

7. Lee HJ, Park JH, Trotter JH, Maher JN, Keenoy KE, Jang YM, et al. Reelin and APP Cooperatively Modulate Dendritic Spine Formation In Vitro and In Vivo. Exp Neurobiol. 2023;32(1):42–55.

8. Chen C, Wei J, Ma X, Xia B, Shakir N, Zhang JK, et al. Disrupted Maturation of Prefrontal Layer 5 Neuronal Circuits in an Alzheimer’s Mouse Model of Amyloid Deposition. Neurosci Bull. 2023;39(6):881–92.

9. Chau DD, Ng LL, Zhai Y, Lau KF. Amyloid precursor protein and its interacting proteins in neurodevelopment. Biochem Soc Trans. 2023;51(4):1647–59.

10. Cvetkovska V, Ge Y, Xu Q, Li S, Zhang P, Craig AM. Neurexin-β Mediates the Synaptogenic Activity of Amyloid Precursor Protein. J Neurosci. 2022;42(48):8936–47.

11. Southam KA, Stennard F, Pavez C, Small DH. Knockout of Amyloid β Protein Precursor (APP) Expression Alters Synaptogenesis, Neurite Branching and Axonal Morphology of Hippocampal Neurons. Neurochem Res. 2019;44(6):1346–55.

12. Lee SH, Kang J, Ho A, Watanabe H, Bolshakov VY, Shen J. APP Family Regulates Neuronal Excitability and Synaptic Plasticity but Not Neuronal Survival. Neuron. 2020;108(4):676–90 e8.

13. Kim JH. Genetics of Alzheimer’s Disease. Dement Neurocogn Disord. 2018;17(4):131–6.

14. Xia D, Watanabe H, Wu B, Lee SH, Li Y, Tsvetkov E, et al. Presenilin-1 knockin mice reveal loss-of-function mechanism for familial Alzheimer’s disease. Neuron. 2015;85(5):967–81.

15. Wolfe MS. Dysfunctional γ-Secretase in Familial Alzheimer’s Disease. Neurochem Res. 2019;44(1):5–11.

16. Esselens C, Sannerud R, Gallardo R, Baert V, Kaden D, Serneels L, et al. Peptides based on the presenilin-APP binding domain inhibit APP processing and Aβ production through interfering with the APP transmembrane domain. Faseb j. 2012;26(9):3765–78.

17. Chavez-Gutierrez L, Bammens L, Benilova I, Vandersteen A, Benurwar M, Borgers M, et al. The mechanism of gamma-Secretase dysfunction in familial Alzheimer disease. EMBO J. 2012;31(10):2261–74.

18. Quintero-Monzon O, Martin MM, Fernandez MA, Cappello CA, Krzysiak AJ, Osenkowski P, et al. Dissociation between the processivity and total activity of gamma-secretase: implications for the mechanism of Alzheimer’s disease-causing presenilin mutations. Biochemistry. 2011;50(42):9023–35.

19. Devkota S, Williams TD, Wolfe MS. Familial Alzheimer’s disease mutations in amyloid protein precursor alter proteolysis by γ-secretase to increase amyloid β-peptides of ≥45 residues. J Biol Chem. 2021;296:100281.

20. Sun L, Zhou R, Yang G, Shi Y. Analysis of 138 pathogenic mutations in presenilin-1 on the in vitro production of Aβ42 and Aβ40 peptides by γ-secretase. Proc Natl Acad Sci U S A. 2017;114(4):E476–e85.

21. Devkota S, Zhou R, Nagarajan V, Maesako M, Do H, Noorani A, et al. Familial Alzheimer mutations stabilize synaptotoxic gamma-secretase-substrate complexes. Cell Rep. 2024;43(2):113761.

22. Cao X, Südhof TC. Dissection of amyloid-beta precursor protein-dependent transcriptional transactivation. J Biol Chem. 2004;279(23):24601–11.

23. Cao X, Südhof TC. A transcriptionally [correction of transcriptively] active complex of APP with Fe65 and histone acetyltransferase Tip60. Science. 2001;293(5527):115–20.

24. Bukhari H, Glotzbach A, Kolbe K, Leonhardt G, Loosse C, Müller T. Small things matter: Implications of APP intracellular domain AICD nuclear signaling in the progression and pathogenesis of Alzheimer’s disease. Prog Neurobiol. 2017;156:189–213.

25. Müller T, Schrötter A, Loosse C, Pfeiffer K, Theiss C, Kauth M, et al. A ternary complex consisting of AICD, FE65, and TIP60 down-regulates Stathmin1. Biochim Biophys Acta. 2013;1834(1):387–94.

26. Müller T, Concannon CG, Ward MW, Walsh CM, Tirniceriu AL, Tribl F, et al. Modulation of gene expression and cytoskeletal dynamics by the amyloid precursor protein intracellular domain (AICD). Mol Biol Cell. 2007;18(1):201–10.

27. Beaver M, Karisetty BC, Zhang H, Bhatnagar A, Armour E, Parmar V, et al. Chromatin and transcriptomic profiling uncover dysregulation of the Tip60 HAT/HDAC2 epigenomic landscape in the neurodegenerative brain. Epigenetics. 2022;17(7):786–807.

28. Sharma S, Sarathlal KC, Taliyan R. Epigenetics in Neurodegenerative Diseases: The Role of Histone Deacetylases. CNS Neurol Disord Drug Targets. 2019;18(1):11–8.

29. Beaver M, Bhatnagar A, Panikker P, Zhang H, Snook R, Parmar V, et al. Disruption of Tip60 HAT mediated neural histone acetylation homeostasis is an early common event in neurodegenerative diseases. Sci Rep. 2020;10(1):18265.

30. Panikker P, Xu SJ, Zhang H, Sarthi J, Beaver M, Sheth A, et al. Restoring Tip60 HAT/HDAC2 Balance in the Neurodegenerative Brain Relieves Epigenetic Transcriptional Repression and Reinstates Cognition. J Neurosci. 2018;38(19):4569–83.

31. Urban I, Kerimoglu C, Sakib MS, Wang H, Benito E, Thaller C, et al. TIP60/KAT5 is required for neuronal viability in hippocampal CA1. Sci Rep. 2019;9(1):16173.

32. Antonell A, Lladó A, Altirriba J, Botta-Orfila T, Balasa M, Fernández M, et al. A preliminary study of the whole-genome expression profile of sporadic and monogenic early-onset Alzheimer’s disease. Neurobiol Aging. 2013;34(7):1772–8.

33. Cary GA, Wiley JC, Gockley J, Keegan S, Amirtha Ganesh SS, Heath L, et al. Genetic and multi-omic risk assessment of Alzheimer’s disease implicates core associated biological domains. Alzheimers Dement (N Y). 2024;10(2):e12461.

34. Oakley H, Cole SL, Logan S, Maus E, Shao P, Craft J, et al. Intraneuronal beta-amyloid aggregates, neurodegeneration, and neuron loss in transgenic mice with five familial Alzheimer’s disease mutations: potential factors in amyloid plaque formation. J Neurosci. 2006;26(40):10129–40.

35. Onos KD, Quinney SK, Jones DR, Masters AR, Pandey R, Keezer KJ, et al. Pharmacokinetic, pharmacodynamic, and transcriptomic analysis of chronic levetiracetam treatment in 5XFAD mice: A MODEL-AD preclinical testing core study. Alzheimers Dement (N Y). 2022;8(1):e12329.

36. Oblak AL, Lin PB, Kotredes KP, Pandey RS, Garceau D, Williams HM, et al. Comprehensive Evaluation of the 5XFAD Mouse Model for Preclinical Testing Applications: A MODEL-AD Study. Front Aging Neurosci. 2021;13:713726.

37. Forner S, Kawauchi S, Balderrama-Gutierrez G, Kramár EA, Matheos DP, Phan J, et al. Systematic phenotyping and characterization of the 5xFAD mouse model of Alzheimer’s disease. Sci Data. 2021;8(1):270.

38. Menden K, Francescatto M, Nyima T, Blauwendraat C, Dhingra A, Castillo-Lizardo M, et al. A multi-omics dataset for the analysis of frontotemporal dementia genetic subtypes. Sci Data. 2023;10(1):849.

39. Yu G, Wang LG, Han Y, He QY. clusterProfiler: an R package for comparing biological themes among gene clusters. Omics. 2012;16(5):284–7.

40. Xie Z, Bailey A, Kuleshov MV, Clarke DJB, Evangelista JE, Jenkins SL, et al. Gene Set Knowledge Discovery with Enrichr. Curr Protoc. 2021;1(3):e90.

41. Sollis E, Mosaku A, Abid A, Buniello A, Cerezo M, Gil L, et al. The NHGRI-EBI GWAS Catalog: knowledgebase and deposition resource. Nucleic Acids Res. 2023;51(D1):D977–D85.

42. Evangelista JE, Xie Z, Marino GB, Nguyen N, Clarke DJB, Ma’ayan A. Enrichr-KG: bridging enrichment analysis across multiple libraries. Nucleic Acids Res. 2023;51(W1):W168–w79.

43. Mishra S, Knupp A, Szabo MP, Williams CA, Kinoshita C, Hailey DW, et al. The Alzheimer’s gene SORL1 is a regulator of endosomal traffic and recycling in human neurons. Cell Mol Life Sci. 2022;79(3):162.

44. Knupp A, Mishra S, Martinez R, Braggin JE, Szabo M, Kinoshita C, et al. Depletion of the AD Risk Gene SORL1 Selectively Impairs Neuronal Endosomal Traffic Independent of Amyloidogenic APP Processing. Cell Rep. 2020;31(9):107719.

45. Yuan SH, Martin J, Elia J, Flippin J, Paramban RI, Hefferan MP, et al. Cell-surface marker signatures for the isolation of neural stem cells, glia and neurons derived from human pluripotent stem cells. PLoS One. 2011;6(3):e17540.

46. Young JE, Fong LK, Frankowski H, Petsko GA, Small SA, Goldstein LSB. Stabilizing the Retromer Complex in a Human Stem Cell Model of Alzheimer’s Disease Reduces TAU Phosphorylation Independently of Amyloid Precursor Protein. Stem Cell Reports. 2018;10(3):1046–58.

47. Israel MA, Yuan SH, Bardy C, Reyna SM, Mu Y, Herrera C, et al. Probing sporadic and familial Alzheimer’s disease using induced pluripotent stem cells. Nature. 2012;482(7384):216–20.

48. Gabitto MI, Travaglini KJ, Rachleff VM, Kaplan ES, Long B, Ariza J, et al. Integrated multimodal cell atlas of Alzheimer’s disease. Nat Neurosci. 2024;27(12):2366–83.

49. Tian R, Abarientos A, Hong J, Hashemi SH, Yan R, Drager N, et al. Genome-wide CRISPRi/a screens in human neurons link lysosomal failure to ferroptosis. Nat Neurosci. 2021;24(7):1020–34.

50. Apostolova LG, Zarow C, Biado K, Hurtz S, Boccardi M, Somme J, et al. Relationship between hippocampal atrophy and neuropathology markers: a 7T MRI validation study of the EADC-ADNI Harmonized Hippocampal Segmentation Protocol. Alzheimers Dement. 2015;11(2):139–50.

51. Fong LK, Yang MM, Dos Santos Chaves R, Reyna SM, Langness VF, Woodruff G, et al. Full-length amyloid precursor protein regulates lipoprotein metabolism and amyloid-β clearance in human astrocytes. J Biol Chem. 2018;293(29):11341–57.

52. Kwart D, Gregg A, Scheckel C, Murphy EA, Paquet D, Duffield M, et al. A Large Panel of Isogenic APP and PSEN1 Mutant Human iPSC Neurons Reveals Shared Endosomal Abnormalities Mediated by APP β-CTFs, Not Aβ. Neuron. 2019;104(2):256–70.e5.

53. Johnson ECB, Carter EK, Dammer EB, Duong DM, Gerasimov ES, Liu Y, et al. Large-scale deep multi-layer analysis of Alzheimer’s disease brain reveals strong proteomic disease-related changes not observed at the RNA level. Nat Neurosci. 2022;25(2):213–25.

54. Rexach JE, Cheng Y, Chen L, Polioudakis D, Lin LC, Mitri V, et al. Cross-disorder and disease-specific pathways in dementia revealed by single-cell genomics. Cell. 2024;187(20):5753–74.e28.

55. Hu Y, Fisher JB, Koprowski S, McAllister D, Kim MS, Lough J. Homozygous disruption of the Tip60 gene causes early embryonic lethality. Dev Dyn. 2009;238(11):2912–21.

56. Tominaga K, Sakashita E, Kasashima K, Kuroiwa K, Nagao Y, Iwamori N, et al. Tip60/KAT5 Histone Acetyltransferase Is Required for Maintenance and Neurogenesis of Embryonic Neural Stem Cells. Int J Mol Sci. 2023;24(3).

57. Humbert J, Salian S, Makrythanasis P, Lemire G, Rousseau J, Ehresmann S, et al. De Novo KAT5 Variants Cause a Syndrome with Recognizable Facial Dysmorphisms, Cerebellar Atrophy, Sleep Disturbance, and Epilepsy. Am J Hum Genet. 2020;107(3):564–74.

58. Shen J, Kelleher RJ, 3rd. The presenilin hypothesis of Alzheimer’s disease: evidence for a loss-of-function pathogenic mechanism. Proc Natl Acad Sci U S A. 2007;104(2):403–9.

59. Wiley JC, Hudson M, Kanning KC, Schecterson LC, Bothwell M. Familial Alzheimer’s disease mutations inhibit gamma-secretase-mediated liberation of beta-amyloid precursor protein carboxy-terminal fragment. J Neurochem. 2005;94(5):1189–201.

60. Mameri A, Côté J. JAZF1: A metabolic actor subunit of the NuA4/TIP60 chromatin modifying complex. Front Cell Dev Biol. 2023;11:1134268.

61. Procida T, Friedrich T, Jack APM, Peritore M, Bönisch C, Eberl HC, et al. JAZF1, A Novel p400/TIP60/NuA4 Complex Member, Regulates H2A.Z Acetylation at Regulatory Regions. Int J Mol Sci. 2021;22(2).

62. Andrews SJ, Renton AE, Fulton-Howard B, Podlesny-Drabiniok A, Marcora E, Goate AM. The complex genetic architecture of Alzheimer’s disease: novel insights and future directions. EBioMedicine. 2023;90:104511.

63. Woodruff G, Young JE, Martinez FJ, Buen F, Gore A, Kinaga J, et al. The presenilin-1 ΔE9 mutation results in reduced γ-secretase activity, but not total loss of PS1 function, in isogenic human stem cells. Cell Rep. 2013;5(4):974–85.

64. Armour EM, Thomas CM, Greco G, Bhatnagar A, Elefant F. Experience-dependent Tip60 nucleocytoplasmic transport is regulated by its NLS/NES sequences for neuroplasticity gene control. Mol Cell Neurosci. 2023;127:103888.

