## Supplemental Figures 1-9 for "Kat5 cKO Biological Domain Signatures Align with Human Alzheimer’s Disease"

### Figure S1

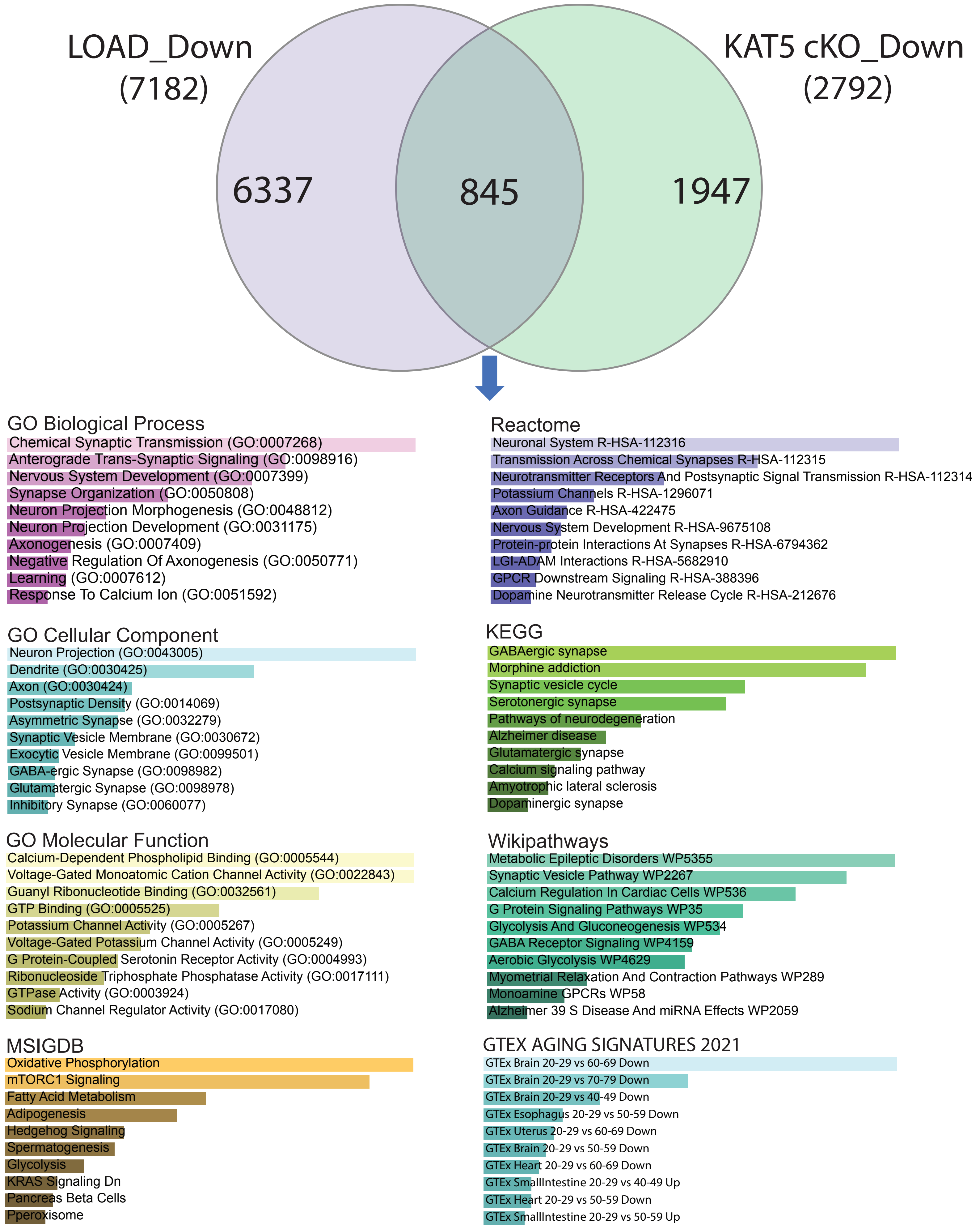
LOAD_Down
(7182)
KAT5 cKO_Down
(2792)
6337
845
1947
GO Biological Process
Chemical Synaptic Transmission (GO:0007268)
Anterograde Trans-Synaptic Signaling (GO:0098916)
Nervous System Development (GO:0007399)
Synapse Organization (GO:0050808)
Neuron Projection Morphogenesis (GO:0048812)
Neuron Projection Development (GO:0031175)
Axonogenesis (GO:0007409)
Negative Regulation Of Axonogenesis (GO:0050771)
Learning (GO:0007612)
Response To Calcium Ion (GO:0051592)
Reactome
Neuronal System R-HSA-112316
Transmission Across Chemical Synapses R-HSA-112315
Neurotransmitter Receptors And Postsynaptic Signal Transmission R-HSA-112314
Potassium Channels R-HSA-1296071
Axon Guidance R-HSA-422475
Nervous System Development R-HSA-9675108
Protein-protein Interactions At Synapses R-HSA-6794362
LGI-ADAM Interactions R-HSA-5682910
GPCR Downstream Signaling R-HSA-388396
Dopamine Neurotransmitter Release Cycle R-HSA-212676
GO Cellular Component
Neuron Projection (GO:0043005)
Dendrite (GO:0030425)
Axon (GO:0030424)
Postsynaptic Density (GO:0014069)
Asymmetric Synapse (GO:0032279)
Synaptic Vesicle Membrane (GO:0030672)
Exocytic Vesicle Membrane (GO:0099501)
GABA-ergic Synapse (GO:0098982)
Glutamatergic Synapse (GO:0098978)
Inhibitory Synapse (GO:0060077)
KEGG
GABAergic synapse
Morphine addiction
Synaptic vesicle cycle
Serotonergic synapse
Pathways of neurodegeneration
Alzheimer disease
Glutamatergic synapse
Calcium signaling pathway
Amyotrophic lateral sclerosis
Dopaminergic synapse
GO Molecular Function
Calcium-Dependent Phospholipid Binding (GO:0005544)
Voltage-Gated Monoatomic Cation Channel Activity (GO:0022843)
Guanyl Ribonucleotide Binding (GO:0032561)
GTP Binding (GO:0005525)
Potassium Channel Activity (GO:0005267)
Voltage-Gated Potassium Channel Activity (GO:0005249)
G Protein-Coupled Serotonin Receptor Activity (GO:0004993)
Ribonucleoside Triphosphate Phosphatase Activity (GO:0017111)
GTPase Activity (GO:0003924)
Sodium Channel Regulator Activity (GO:0017080)
Wikipathways
Metabolic Epileptic Disorders WP5355
Synaptic Vesicle Pathway WP2267
Calcium Regulation In Cardiac Cells WP536
G Protein Signaling Pathways WP35
Glycolysis And Gluconeogenesis WP534
GABA Receptor Signaling WP4159
Aerobic Glycolysis WP4629
Myometrial Relaxation And Contraction Pathways WP289
Monoamine GPCRs WP58
Alzheimer 39 S Disease And miRNA Effects WP2059
MSIGDB
Oxidative Phosphorylation
mTORC1 Signaling
Fatty Acid Metabolism
Adipogenesis
Hedgehog Signaling
Spermatogenesis
Glycolysis
KRAS Signaling Dn
Pancreas Beta Cells
Pperoxisome
GTEX AGING SIGNATURES 2021
GTEx Brain 20-29 vs 60-69 Down
GTEx Brain 20-29 vs 70-79 Down
GTEx Brain 20-29 vs 40-49 Down
GTEx Esophagus 20-29 vs 50-59 Down
GTEx Uterus 20-29 vs 60-69 Down
GTEx Brain 20-29 vs 50-59 Down
GTEx Heart 20-29 vs 60-69 Down
GTEx SmallIntestine 20-29 vs 40-49 Up
GTEx Heart 20-29 vs 50-59 Down
GTEx SmallIntestine 20-29 vs 50-59 Up

Supplementary Figure 1. Convergence of Down-Regulated Genes in Kat5 cKO and LOAD Across GO and Pathway Models.

Drawing from our recent work harmonizing over 1700 postmortem brain bank transcriptomic profiles, we identify a set of 7182 significantly down-regulated genes within LOAD and compare those to the 2792 unique down-regulated Kat5 cKO genes ($p<0.05$, 0.2 differential expression cut-off). There are 845 genes shared between the two groups that are employed in enrichment studies of GO term enrichment within each of the primary tiers of GO (biological process, cellular component, and molecular function). Pathway enrichment analysis was performed using the same 845 genes across Reactome, KEGG, and WikiPathways. The 845 shared genes overlap the GTEX signatures for brain aging (brain 20-29 vs 60-69; 20-29 vs 70-79) most highly, consistent with the alignment of the Kat5 cKO model with human brain aging. All colored bars are significant at $p<0.05$, with shorter bars denoting decreased p-values and darker colors representing decreasing enrichment scores. All values are available in Supplementary Table 2.

Figure S2

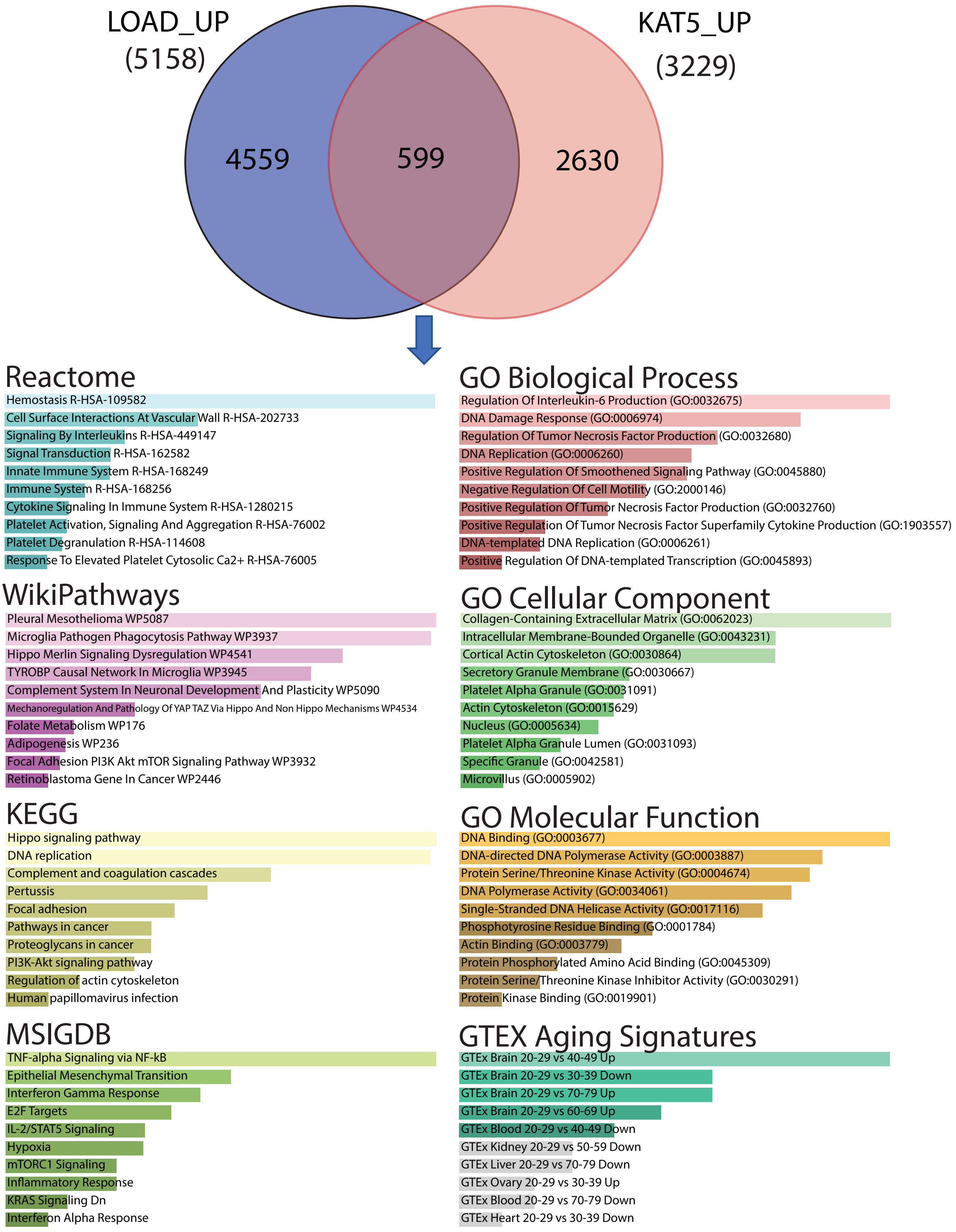
LOAD_UP
(5158)
KAT5_UP
(3229)
4559
599
2630
Reactome
Hemostasis R-HSA-109582
Cell Surface Interactions At Vascular Wall R-HSA-202733
Signaling By Interleukins R-HSA-449147
Signal Transduction R-HSA-162582
Innate Immune System R-HSA-168249
Immune System R-HSA-168256
Cytokine Signaling In Immune System R-HSA-1280215
Platelet Activation, Signaling And Aggregation R-HSA-76002
Platelet Degranulation R-HSA-114608
Response To Elevated Platelet Cytosolic Ca2+ R-HSA-76005
GO Biological Process
Regulation Of Interleukin-6 Production (GO:0032675)
DNA Damage Response (GO:0006974)
Regulation Of Tumor Necrosis Factor Production (GO:0032680)
DNA Replication (GO:0006260)
Positive Regulation Of Smoothened Signaling Pathway (GO:0045880)
Negative Regulation Of Cell Motility (GO:2000146)
Positive Regulation Of Tumor Necrosis Factor Production (GO:0032760)
Positive Regulation Of Tumor Necrosis Factor Superfamily Cytokine Production (GO:1903557)
DNA-templated DNA Replication (GO:0006261)
Positive Regulation Of DNA-templated Transcription (GO:0045893)
WikiPathways
Pleural Mesothelioma WP5087
Microglia Pathogen Phagocytosis Pathway WP3937
Hippo Merlin Signaling Dysregulation WP4541
TYROBP Causal Network In Microglia WP3945
Complement System In Neuronal Development And Plasticity WP5090
Mechanoregulation And Pathology Of YAP TAZ Via Hippo And Non Hippo Mechanisms WP4534
Folate Metabolism WP176
Adipogenesis WP236
Focal Adhesion PI3K Akt mTOR Signaling Pathway WP3932
Retinoblastoma Gene In Cancer WP2446
GO Cellular Component
Collagen-Containing Extracellular Matrix (GO:0062023)
Intracellular Membrane-Bounded Organelle (GO:0043231)
Cortical Actin Cytoskeleton (GO:0030864)
Secretory Granule Membrane (GO:0030667)
Platelet Alpha Granule (GO:0031091)
Actin Cytoskeleton (GO:0015629)
Nucleus (GO:0005634)
Platelet Alpha Granule Lumen (GO:0031093)
Specific Granule (GO:0042581)
Microvillus (GO:0005902)
KEGG
Hippo signaling pathway
DNA replication
Complement and coagulation cascades
Pertussis
Focal adhesion
Pathways in cancer
Proteoglycans in cancer
PI3K-Akt signaling pathway
Regulation of actin cytoskeleton
Human papillomavirus infection
GO Molecular Function
DNA Binding (GO:0003677)
DNA-directed DNA Polymerase Activity (GO:0003887)
Protein Serine/Threonine Kinase Activity (GO:0004674)
DNA Polymerase Activity (GO:0034061)
Single-Stranded DNA Helicase Activity (GO:0017116)
Phosphotyrosine Residue Binding (GO:0001784)
Actin Binding (GO:0003779)
Protein Phosphorylated Amino Acid Binding (GO:0045309)
Protein Serine/Threonine Kinase Inhibitor Activity (GO:0030291)
Protein Kinase Binding (GO:0019901)
MSIGDB
TNF-alpha Signaling via NF-kB
Epithelial Mesenchymal Transition
Interferon Gamma Response
E2F Targets
IL-2/STAT5 Signaling
Hypoxia
mTORC1 Signaling
Inflammatory Response
KRAS Signaling Dn
Interferon Alpha Response
GTEX Aging Signatures
GTEx Brain 20-29 vs 40-49 Up
GTEx Brain 20-29 vs 30-39 Down
GTEx Brain 20-29 vs 70-79 Up
GTEx Brain 20-29 vs 60-69 Up
GTEx Blood 20-29 vs 40-49 Down
GTEx Kidney 20-29 vs 50-59 Down
GTEx Liver 20-29 vs 70-79 Down
GTEx Ovary 20-29 vs 30-39 Up
GTEx Blood 20-29 vs 70-79 Down
GTEx Heart 20-29 vs 30-39 Down

Figure S2. Convergence of Up-regulated Genes in Kat5 cKO and LOAD Across GO and Pathway Models.

The TREAT-AD Consortium Bioinformatics Team harmonized over 1700 transcriptomic profiles from harmonized postmortem brain banks. We employed the up-regulated differentially expressed genes (DEG, total 5158) ($p<0.05$) in a comparison with Kat5 cKO up-regulated DEGs (0.2 differential expression cut-off, $p<0.05$, total 3229). The Venn diagram demonstrates the intersection of these two genesets with 599 genes shared between the two datasets. Gene set enrichment analysis was performed with the 599 overlapping genes with all three tiers of GO, select pathway models (Reactome, KEGG, WikiPathways), and the molecular signatures associated with MSIGDB and GTEX aging profiles. All colored bars are significant at $p<0.05$, with shorter bars denoting decreased p-values and darker colors representing decreasing enrichment scores. All values are available in Supplementary Table 3.

### Figure S3

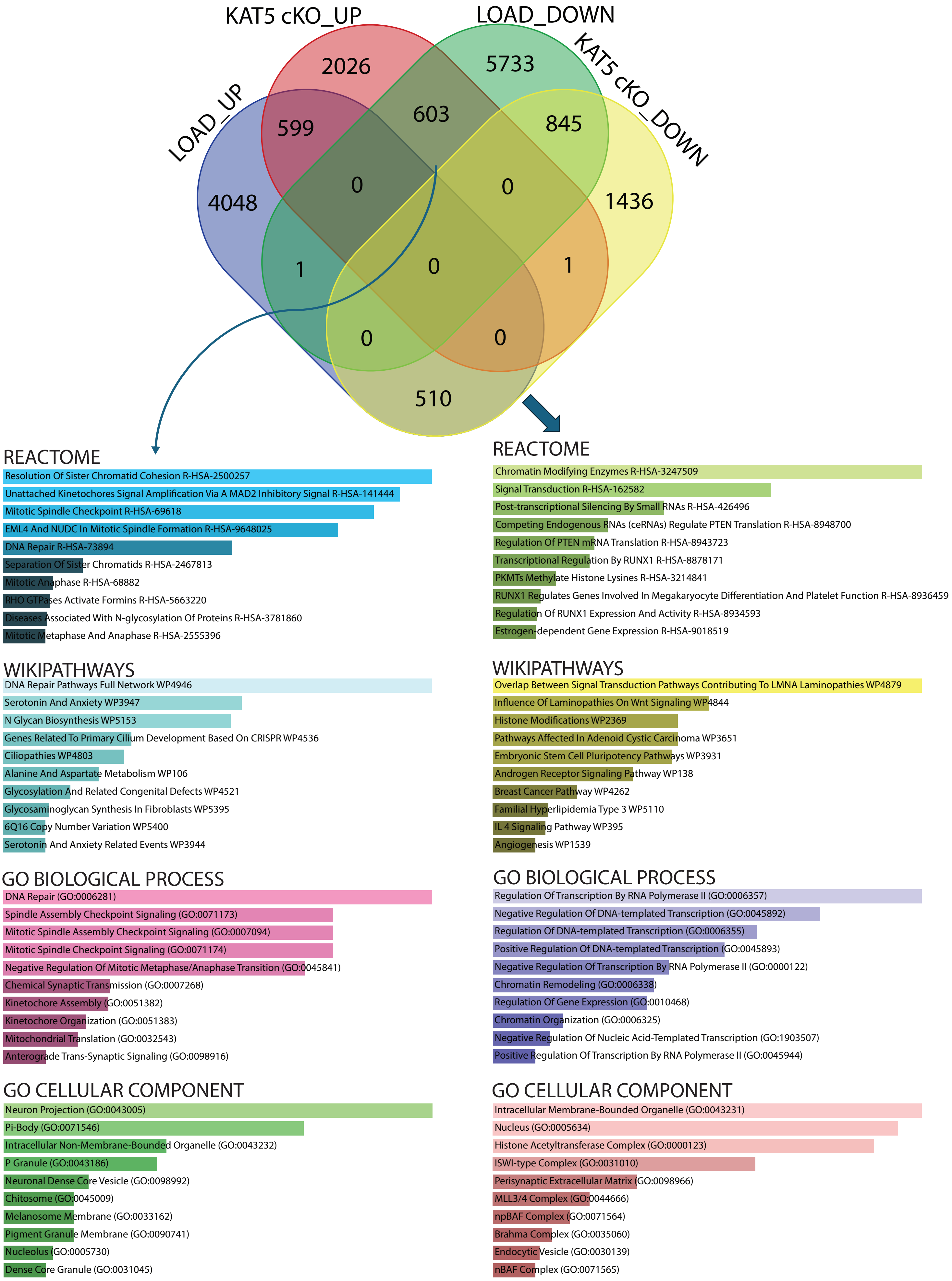
KAT5 cKO_UP
LOAD_DOWN
LOAD_UP
KAT5 cKO_DOWN
2026
5733
599
603
845
4048
0
0
1436
1
0
1
0
0
510
REACTOME
Resolution Of Sister Chromatid Cohesion R-HSA-2500257
Unattached Kinetochores Signal Amplification Via A MAD2 Inhibitory Signal R-HSA-141444
Mitotic Spindle Checkpoint R-HSA-69618
EML4 And NUDC In Mitotic Spindle Formation R-HSA-9648025
DNA Repair R-HSA-73894
Separation Of Sister Chromatids R-HSA-2467813
Mitotic Anaphase R-HSA-68882
RHO GTPases Activate Formins R-HSA-5663220
Diseases Associated With N-glycosylation Of Proteins R-HSA-3781860
Mitotic Metaphase And Anaphase R-HSA-2555396
WIKIPATHWAYS
DNA Repair Pathways Full Network WP4946
Serotonin And Anxiety WP3947
N Glycan Biosynthesis WP5153
Genes Related To Primary Cilium Development Based On CRISPR WP4536
Ciliopathies WP4803
Alanine And Aspartate Metabolism WP106
Glycosylation And Related Congenital Defects WP4521
Glycosaminoglycan Synthesis In Fibroblasts WP5395
6Q16 Copy Number Variation WP5400
Serotonin And Anxiety Related Events WP3944
GO BIOLOGICAL PROCESS
DNA Repair (GO:0006281)
Spindle Assembly Checkpoint Signaling (GO:0071173)
Mitotic Spindle Assembly Checkpoint Signaling (GO:0007094)
Mitotic Spindle Checkpoint Signaling (GO:0071174)
Negative Regulation Of Mitotic Metaphase/Anaphase Transition (GO:0045841)
Chemical Synaptic Transmission (GO:0007268)
Kinetochore Assembly (GO:0051382)
Kinetochore Organization (GO:0051383)
Mitochondrial Translation (GO:0032543)
Anterograde Trans-Synaptic Signaling (GO:0098916)
GO CELLULAR COMPONENT
Neuron Projection (GO:0043005)
Pi-Body (GO:0071546)
Intracellular Non-Membrane-Bounded Organelle (GO:0043232)
P Granule (GO:0043186)
Neuronal Dense Core Vesicle (GO:0098992)
Chitosome (GO:0045009)
Melanosome Membrane (GO:0033162)
Pigment Granule Membrane (GO:0090741)
Nucleolus (GO:0005730)
Dense Core Granule (GO:0031045)
REACTOME
Chromatin Modifying Enzymes R-HSA-3247509
Signal Transduction R-HSA-162582
Post-transcriptional Silencing By Small RNAs R-HSA-426496
Competing Endogenous RNAs (ceRNAs) Regulate PTEN Translation R-HSA-8948700
Regulation Of PTEN mRNA Translation R-HSA-8943723
Transcriptional Regulation By RUNX1 R-HSA-8878171
PKMTs Methylate Histone Lysines R-HSA-3214841
RUNX1 Regulates Genes Involved In Megakaryocyte Differentiation And Platelet Function R-HSA-8936459
Regulation Of RUNX1 Expression And Activity R-HSA-8934593
Estrogen-dependent Gene Expression R-HSA-9018519
WIKIPATHWAYS
Overlap Between Signal Transduction Pathways Contributing To LMNA Laminopathies WP4879
Influence Of Laminopathies On Wnt Signaling WP4844
Histone Modifications WP2369
Pathways Affected In Adenoid Cystic Carcinoma WP3651
Embryonic Stem Cell Pluripotency Pathways WP3931
Androgen Receptor Signaling Pathway WP138
Breast Cancer Pathway WP4262
Familial Hyperlipidemia Type 3 WP5110
IL 4 Signaling Pathway WP395
Angiogenesis WP1539
GO BIOLOGICAL PROCESS
Regulation Of Transcription By RNA Polymerase II (GO:0006357)
Negative Regulation Of DNA-templated Transcription (GO:0045892)
Regulation Of DNA-templated Transcription (GO:0006355)
Positive Regulation Of DNA-templated Transcription (GO:0045893)
Negative Regulation Of Transcription By RNA Polymerase II (GO:0000122)
Chromatin Remodeling (GO:0006338)
Regulation Of Gene Expression (GO:0010468)
Chromatin Organization (GO:0006325)
Negative Regulation Of Nucleic Acid-Templated Transcription (GO:1903507)
Positive Regulation Of Transcription By RNA Polymerase II (GO:0045944)
GO CELLULAR COMPONENT
Intracellular Membrane-Bounded Organelle (GO:0043231)
Nucleus (GO:0005634)
Histone Acetyltransferase Complex (GO:0000123)
ISWI-type Complex (GO:0031010)
Perisynaptic Extracellular Matrix (GO:0098966)
MLL3/4 Complex (GO:0044666)
npBAF Complex (GO:0071564)
Brahma Complex (GO:0035060)
Endocytic Vesicle (GO:0030139)
nBAF Complex (GO:0071565)

Figure Legend S3. Convergence Patterns of Up- and Down-regulated Genes from LOAD and Kat5 cKO.

The LOAD DEGs identified and compared in Supplementary Figures 1 and 2 are employed here in association with the Kat5 cKO DEGs compared in the same figures. The across valence comparison for LOAD and Kat5 cKO mouse model demonstrate that, in addition to the tandem up or down DEGs, there are also substantial gene sets that are cross-regulated within LOAD and Kat5 cKO. The four-way Venn diagram at the top of the figure shows are intersections, including those already demonstrated in the previous figures, and additionally show a set of 603 genes that are up-regulated in Kat5 cKO and down-regulated in LOAD. Conversely, there is a set of 510 genes that are up-regulated in LOAD and down-regulated in Kat5 cKO. Both cross-regulated gene sets are examined with gene set enrichment analysis using Reactome and WikiPathways pathway models and GO biological process and GO Cellular Component. The up-regulated Kat5 cKO genes down-regulated in LOAD prominently reflect DNA repair and cell cycle processes while the down-regulated Kat5 cKO and up-regulated LOAD genes represent largely transcriptional or chromatin modifying processes. All colored bars are significant at $p<0.05$, with shorter bars denoting decreased p-values and darker colors representing decreasing enrichment scores.

### Figure S4

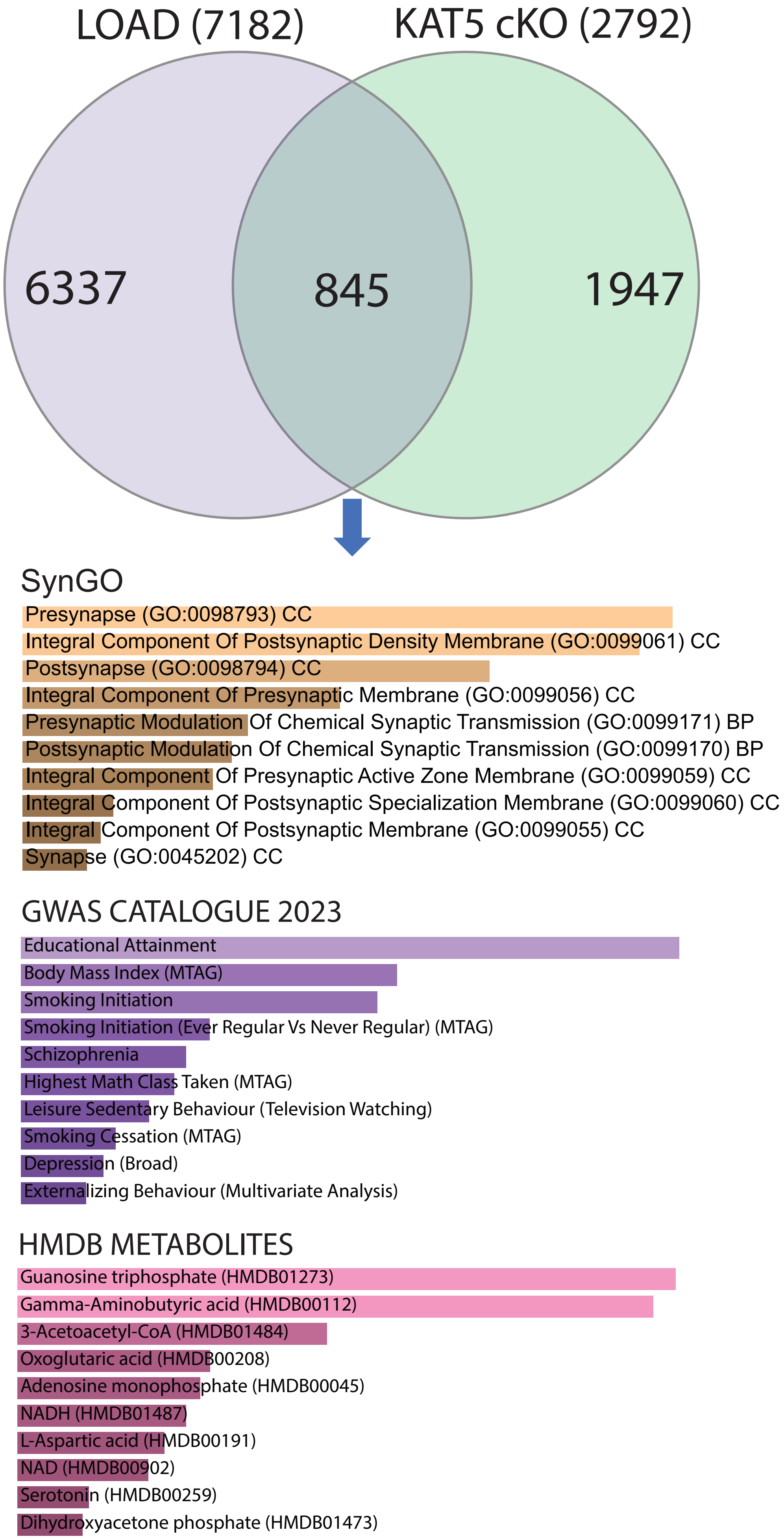

LOAD (7182)
KAT5 cKO (2792)
6337
845
1947
SynGO
Presynapse (GO:0098793) CC
Integral Component Of Postsynaptic Density Membrane (GO:0099061) CC
Postsynapse (GO:0098794) CC
Integral Component Of Presynaptic Membrane (GO:0099056) CC
Presynaptic Modulation Of Chemical Synaptic Transmission (GO:0099171) BP
Postsynaptic Modulation Of Chemical Synaptic Transmission (GO:0099170) BP
Integral Component Of Presynaptic Active Zone Membrane (GO:0099059) CC
Integral Component Of Postsynaptic Specialization Membrane (GO:0099060) CC
Integral Component Of Postsynaptic Membrane (GO:0099055) CC
Synapse (GO:0045202) CC
GWAS CATALOGUE 2023
Educational Attainment
Body Mass Index (MTAG)
Smoking Initiation
Smoking Initiation (Ever Regular Vs Never Regular) (MTAG)
Schizophrenia
Highest Math Class Taken (MTAG)
Leisure Sedentary Behaviour (Television Watching)
Smoking Cessation (MTAG)
Depression (Broad)
Externalizing Behaviour (Multivariate Analysis)
HMDB METABOLITES
Guanosine triphosphate (HMDB01273)
Gamma-Aminobutyric acid (HMDB00112)
3-Acetoacetyl-CoA (HMDB01484)
Oxoglutaric acid (HMDB00208)
Adenosine monophosphate (HMDB00045)
NADH (HMDB01487)
L-Aspartic acid (HMDB00191)
NAD (HMDB00902)
Serotonin (HMDB00259)
Dihydroxyacetone phosphate (HMDB01473)

Figure Legend S4. Additional Analyses of the intersection of the LOAD and Kat5 cKO down-regulated gene sets.

The down-regulated LOAD genes from the TREAT-AD bioinformatics pipeline analysis are employed in conjunction with the 0.2 or greater down-regulated Kat5 cKO genes, resulting in the previously discussed 845 genes common to both gene sets. These are further analysis by gene set enrichment analysis against the GTEX aging signatures, the GWAS Catalogue gene variant-trait linkages, and the human metabolite database (HMDB). The GWAS catalogue shows the greatest overlap with educational attainment, suggesting a link between the Kat5 cKO regulated genes, those that decrement in AD, and genes enhanced by educational achievement. Interestingly, the HMDB signatures that are most highly shared represent elements of core metabolism and neurotransmitters synthesis, such as GABA and Serotonin. SynGO enrichments were included to show synaptically filtered process (SynGO). All colored bars are significant at $p<0.05$, with shorter bars denoting decreased p-values and darker colors representing decreasing enrichment scores. Exact enrichment values and scores are contained within Supplementary table 2.

### Figure S5

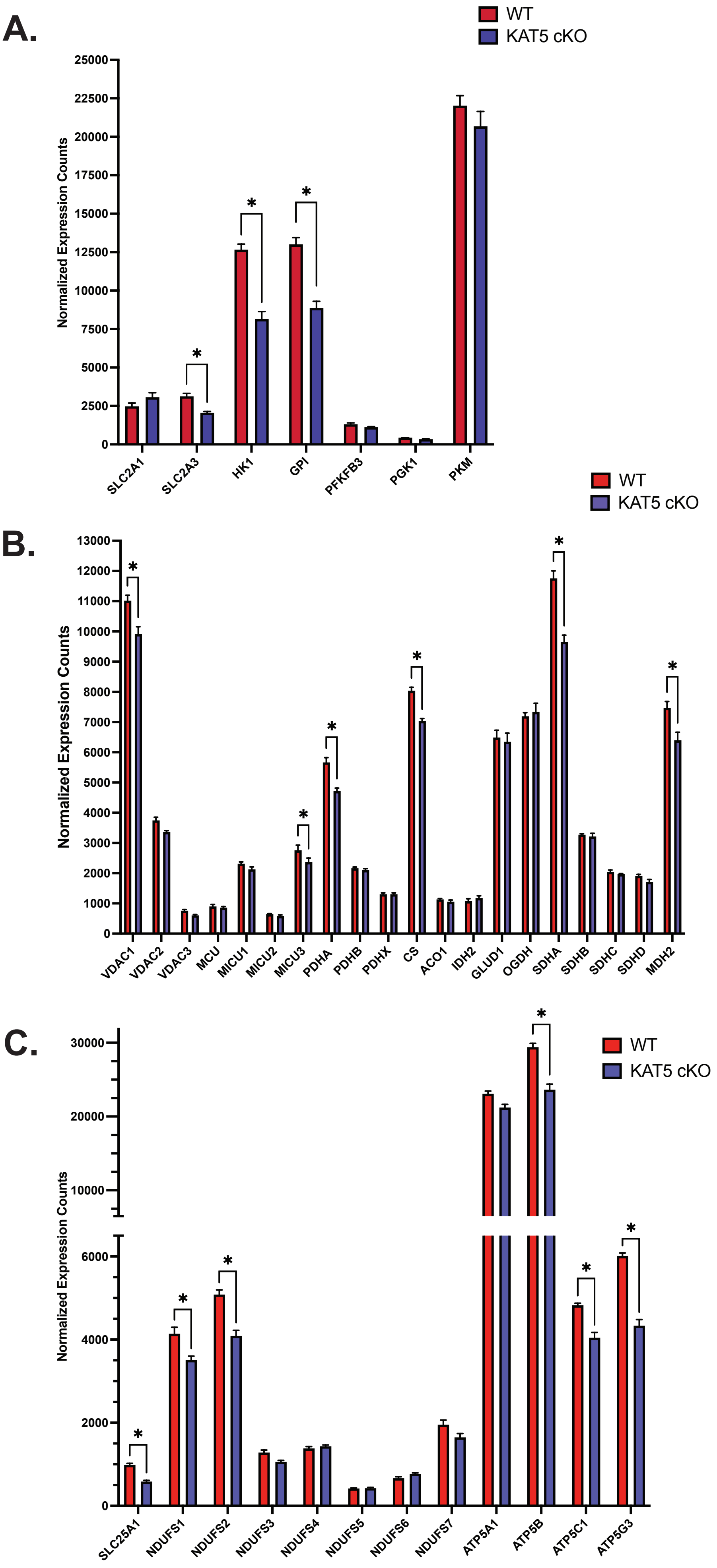

A.
WT
KAT5 cKO
Normalized Expression Counts
25000
22500
20000
17500
15000
12500
10000
7500
5000
2500
0
*
SLC2A1
SLC2A3
HK1
GPI
PFKFB3
PGK1
PKM
B.
WT
KAT5 cKO
Normalized Expression Counts
13000
12000
11000
10000
9000
8000
7000
6000
5000
4000
3000
2000
1000
0
VDAC1
VDAC2
VDAC3
MCU
MICU1
MICU2
MICU3
PDHA
PDHB
PDHX
CS
ACO1
IDH2
GLUD1
OGDH
SDHA
SDHB
SDHC
SDHD
MDH2
C.
WT
KAT5 cKO
Normalized Expression Counts
30000
20000
10000
6000
4000
2000
0
SLC25A1
NDUFS1
NDUFS2
NDUFS3
NDUFS4
NDUFS5
NDUFS6
NDUFS7
ATP5A1
ATP5B
ATP5C1
ATP5G3

Figure Legend S5. Analysis of the glycolysis, tricarboxylic acid (TCA) cycle, and electron transport chain (ETC) genes in the Kat5 cKO vs control.

The six animals per genotype employed in the Kat5 DEG comparison study were investigated for specific differences in defined genes associated with stages of metabolism. The core enzymes in the glycolytic pathway are compared in (A) including an examination of glucose uptake from the neurovasculature by astrocytes, the neuronal glucose uptake from the astrocytic endfeet, and the processing to pyruvate through the core glycolytic enzymes. The biological process analyzed is visually depicted to the right side of the graphs. The neuron specific glucose transporter SLC2A3 is significantly down-regulated within the Kat5 cKO model compared to wild-type. Additionally, HK1 and GPI, the initial enzymatic steps within glycolysis, are also downregulated. The enzymes involved in pyruvate transport and conversion into acetyl-CoA and subsequent processing through the TCA cycle is depicted in (B) with graphical representation of both calcium import factors and the TCA cycle represented to the right. There are isolated decrements in both transport and TCA cycle genes—specifically PDHA, CS, SDHA and MDH2—within the Kat5 cKO model. The transcript levels for the core enzyme involved in conversion of NADH and FADH2 into ATP through the ETC are examined in (C). There are small decreases distributed across complex I and complex V observed, as noted by the asterix ($padj<0.05$).

Figure S6

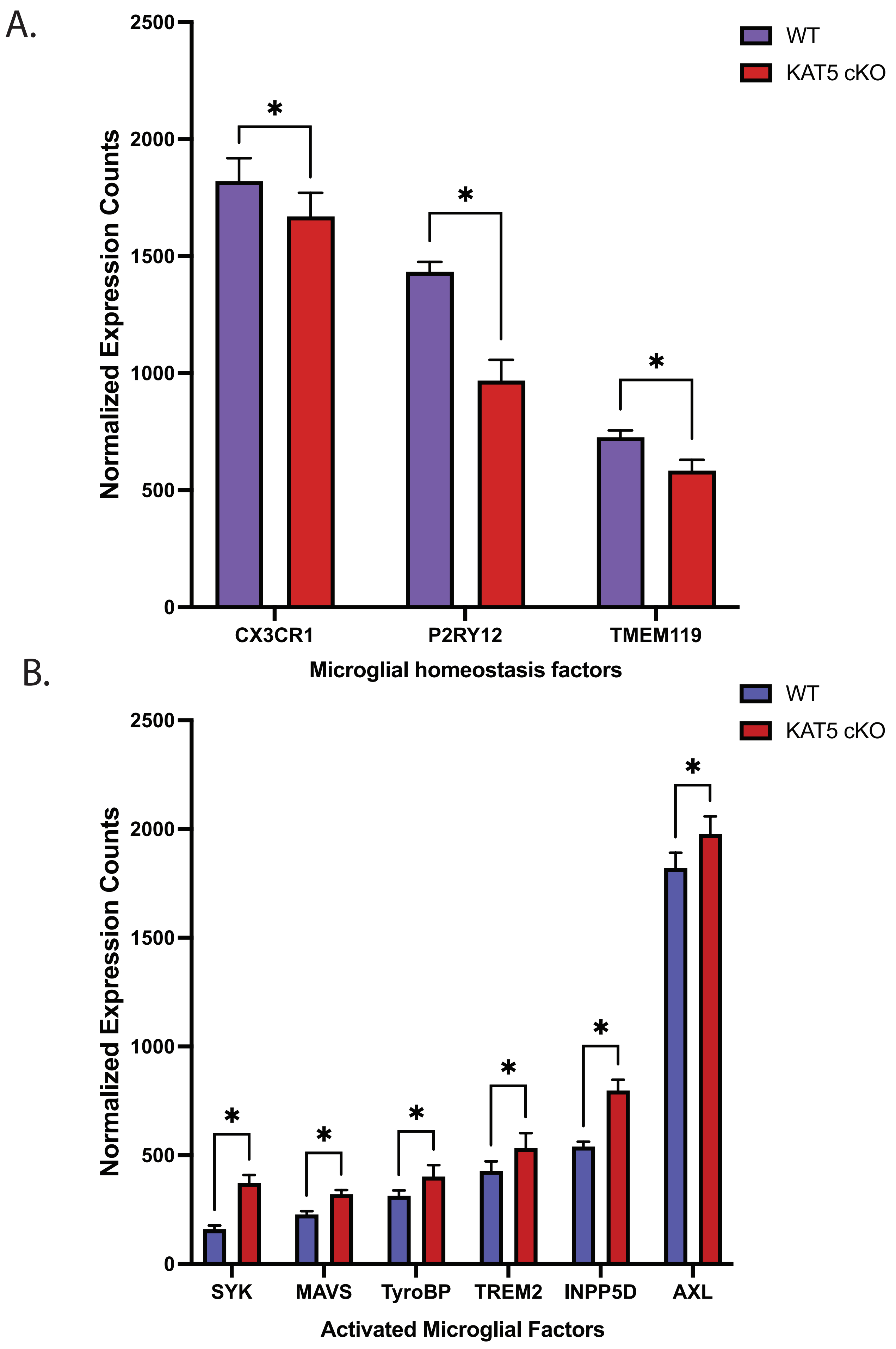
A.
Normalized Expression Counts
2500
2000
1500
1000
500
0
WT
KAT5 cKO
*
*
*
CX3CR1
P2RY12
TMEM119
Microglial homeostasis factors
B.
WT
KAT5 cKO
Normalized Expression Counts
2500
2000
1500
1000
500
0
*
*
*
*
*
*
SYK
MAVS
TyroBP
TREM2
INPP5D
AXL
Activated Microglial Factors

Figure Legend S6. Kat5 cKO downregulates homeostatic microglial factors and upregulates activated microglial markers.

The phenotype of the Kat5 cKO involves progressive reactive gliosis, so we examined the levels of known homeostatic microglial markers (A) and activated microglial markers (B) within the transcriptomic signature occurring prior to pathological encroachment through a comparison of the six Kat5 cKO mice compared to the wild-type controls. The Kat5 cKO mice had small but statistically significant decreases in CX3CR1, P2RY12 and TMEM119 as shown in (A) with the asterix representing significance at the padj<0.05. Conversely, the activated markers SYK, MAVS, TYROBP, TREM2, INPP5D and AXL are all upregulated by small but significant margins (B), with the asterix bar comparison denoting padj<0.05. The overall scope of the change is small, the transcriptomic signature was obtained from mice two weeks after induced gene ablation.

A. **KAT5 Expression change across disease pseudo-progression - Subclass Resolution**

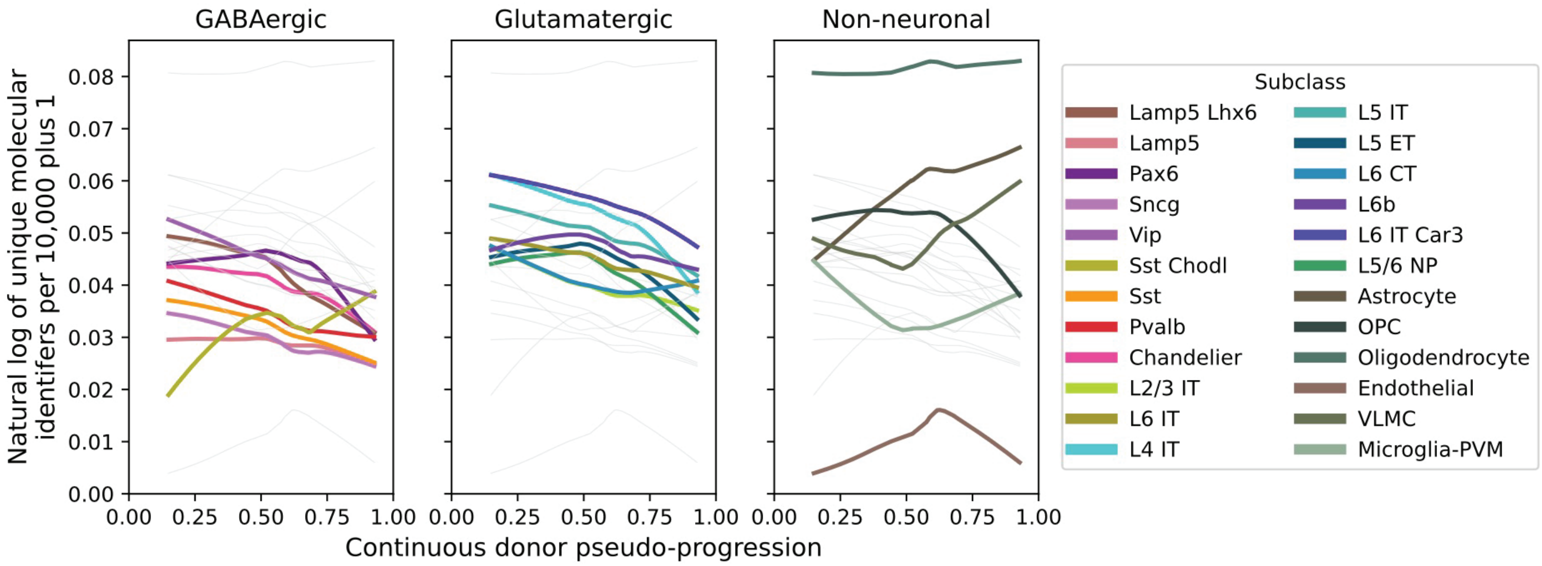


B.

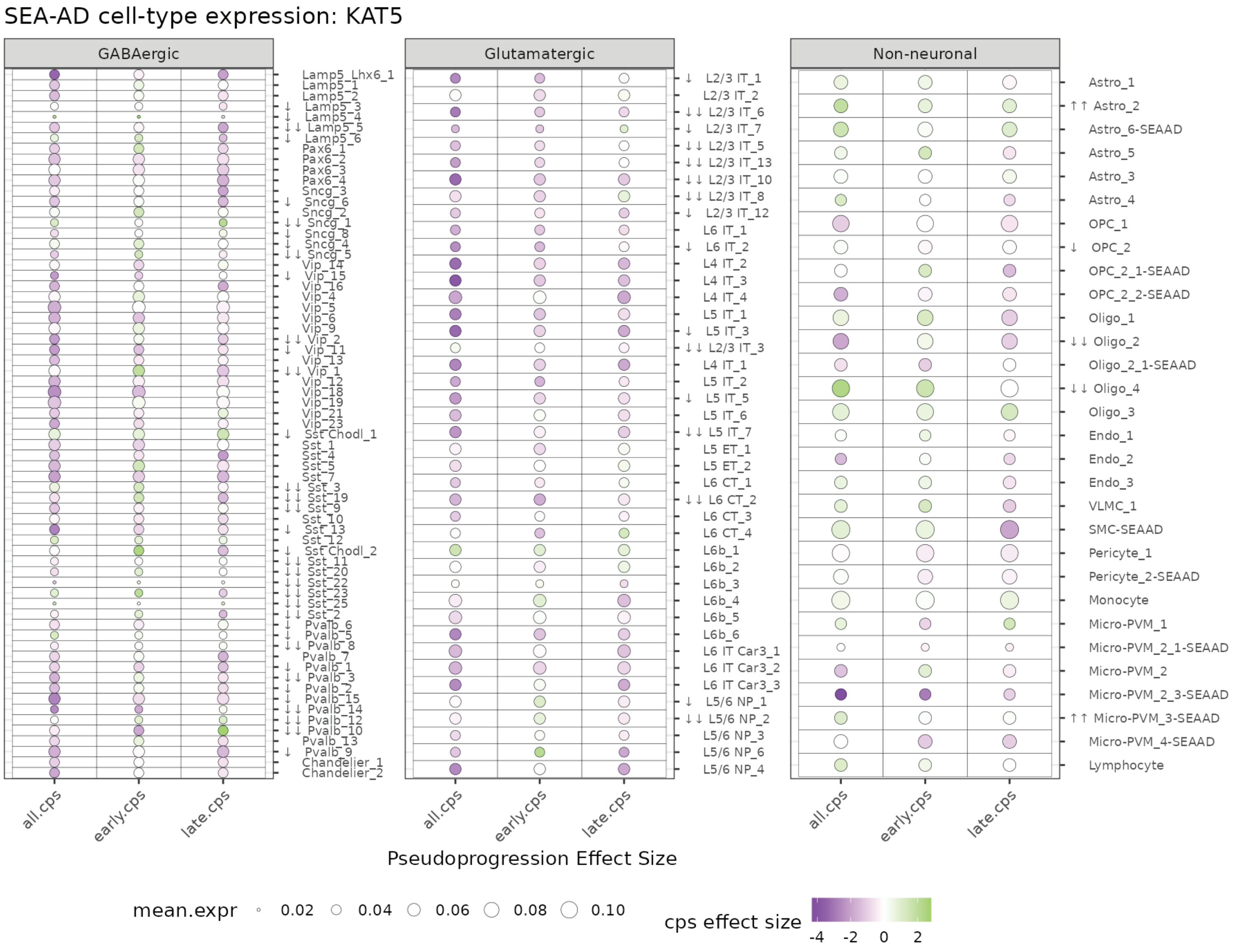

Figure 7. Gamma-Secretase Components Dysregulated in Neurons in SEA-AD LOAD Study

The Seattle Alzheimer's Disease (SEA-AD) study performed expression analysis across AD progression as modeled by the continuous pseudo-progression score (CPS) within all identified cellular subtypes within the human brain taxonomy. The gamma-secretase components are shown at two levels of cellular resolution with column on the left depicting subtype (more general) cell types and the column on data on the right hand side showing all available cellular subtypes. Both views divide the cellular lineages into inhibitory GABAergic neurons, excitatory glutamatergic neurons, and non-neuronal lineages (glial and endothelial subtypes). The representations on the left quantitate expression across disease based upon normalized expression levels and on right the expression is heat-mapped based on CPS effect size with down-regulation represented in purple and upregulation represented in green and the size of the circle representing mean expression level. The homologues of presenilin PSEN1(A) and PSEN2(B) levels across disease are shown in the top two rows. The remains three components of the heterotetrameric gamma-secretase complex are shown below (C) PSENEN, (D) APH1B, and (E) NCSTN. At the low (left) cellular specificity and high cellular specificity (right) representations, there are strong downward regulation of PSEN2 and APH1B within both inhibitory and excitatory neurons.

### Figure S8

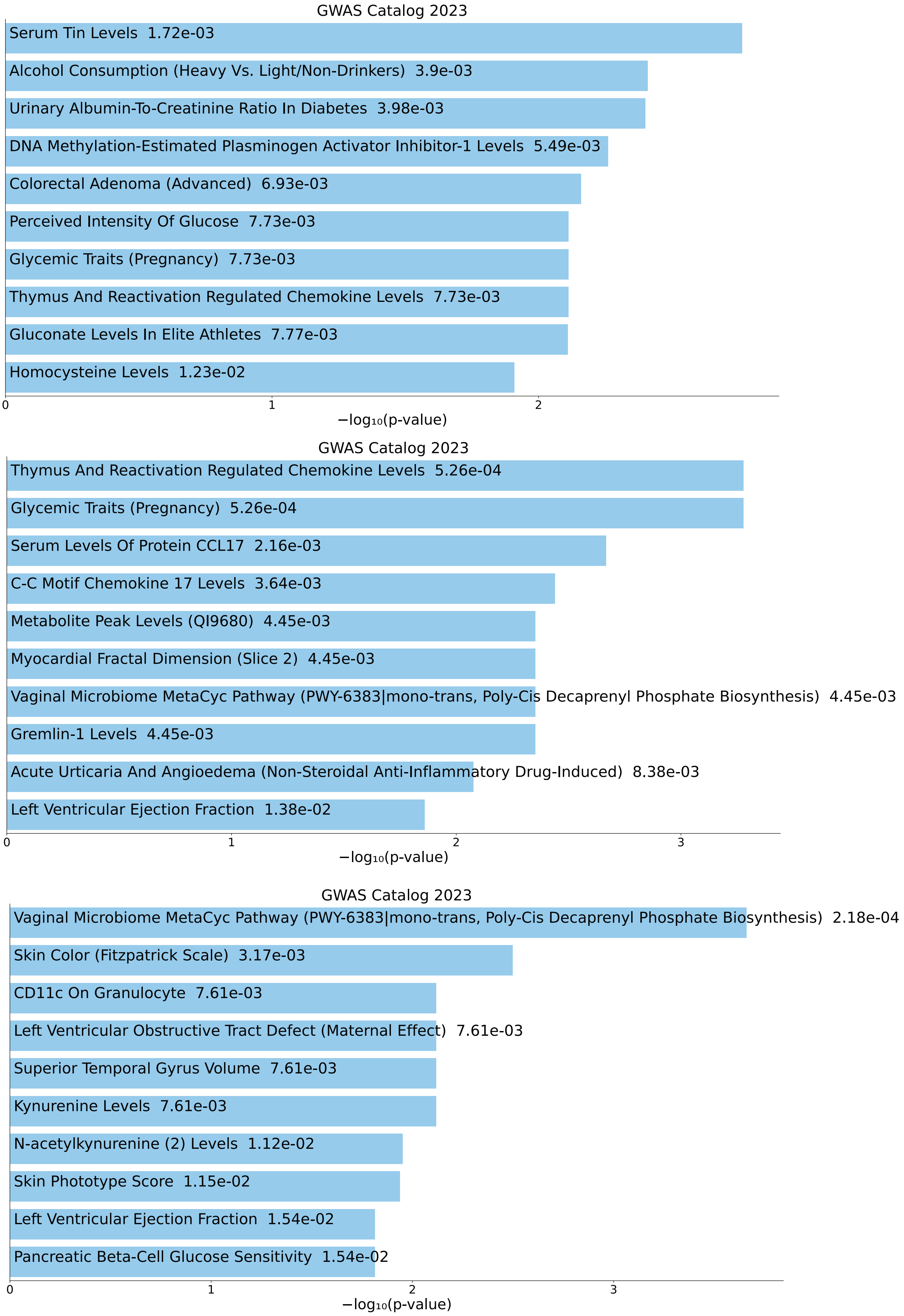
GWAS Catalog 2023
Serum Tin Levels 1.72e-03
Alcohol Consumption (Heavy Vs. Light/Non-Drinkers) 3.9e-03
Urinary Albumin-To-Creatinine Ratio In Diabetes 3.98e-03
DNA Methylation-Estimated Plasminogen Activator Inhibitor-1 Levels 5.49e-03
Colorectal Adenoma (Advanced) 6.93e-03
Perceived Intensity Of Glucose 7.73e-03
Glycemic Traits (Pregnancy) 7.73e-03
Thymus And Reactivation Regulated Chemokine Levels 7.73e-03
Gluconate Levels In Elite Athletes 7.77e-03
Homocysteine Levels 1.23e-02
0
1
2
−log10(p-value)
GWAS Catalog 2023
Thymus And Reactivation Regulated Chemokine Levels 5.26e-04
Glycemic Traits (Pregnancy) 5.26e-04
Serum Levels Of Protein CCL17 2.16e-03
C-C Motif Chemokine 17 Levels 3.64e-03
Metabolite Peak Levels (QI9680) 4.45e-03
Myocardial Fractal Dimension (Slice 2) 4.45e-03
Vaginal Microbiome MetaCyc Pathway (PWY-6383|mono-trans, Poly-Cis Decaprenyl Phosphate Biosynthesis) 4.45e-03
Gremlin-1 Levels 4.45e-03
Acute Urticaria And Angioedema (Non-Steroidal Anti-Inflammatory Drug-Induced) 8.38e-03
Left Ventricular Ejection Fraction 1.38e-02
0
1
2
3
−log10(p-value)
GWAS Catalog 2023
Vaginal Microbiome MetaCyc Pathway (PWY-6383|mono-trans, Poly-Cis Decaprenyl Phosphate Biosynthesis) 2.18e-04
Skin Color (Fitzpatrick Scale) 3.17e-03
CD11c On Granulocyte 7.61e-03
Left Ventricular Obstructive Tract Defect (Maternal Effect) 7.61e-03
Superior Temporal Gyrus Volume 7.61e-03
Kynurenine Levels 7.61e-03
N-acetylkynurenine (2) Levels 1.12e-02
Skin Phototype Score 1.15e-02
Left Ventricular Ejection Fraction 1.54e-02
Pancreatic Beta-Cell Glucose Sensitivity 1.54e-02
0
1
2
3
−log10(p-value)

Figure Legend S8. The GWAS Catalogue gene variant-trait linkages within the upregulated geneset of the Kat5 cKO mice.

The comparison of the 1.2 fold (3229 genes), 1.5 fold (1592), and 2 fold (568) upregulated genes statistically significant at $p<0.05$ post FDR correction were examined for gene variant associations with identified traits within the GWAS Catalogue studies. This comparison is the converse analysis as performed in Figure 9 examining the downregulated genes. The overall significance levels are lower in the upregulated genes than observed in the downregulated gene set, with less evidence of clear specific genetic associations. There does appear to be an association with some cardiac and vascular function, as homocysteine levels are observed within the 1.2 fold upregulated gene set trait associations, myocardial fractal dimension, acute urticaria and angioedema and left ventricular ejection fraction are enriched within the 1.5 fold upregulated gene set and left ventricular obstructive tract defect and left ventricular ejection fraction are weakly associated with the 2.0 fold enriched set. Overall, the patterns of gene variant-trait association in the upregulated genes appear less identifiable and weaker than in the downregulated gene set.

Supp Figure 9

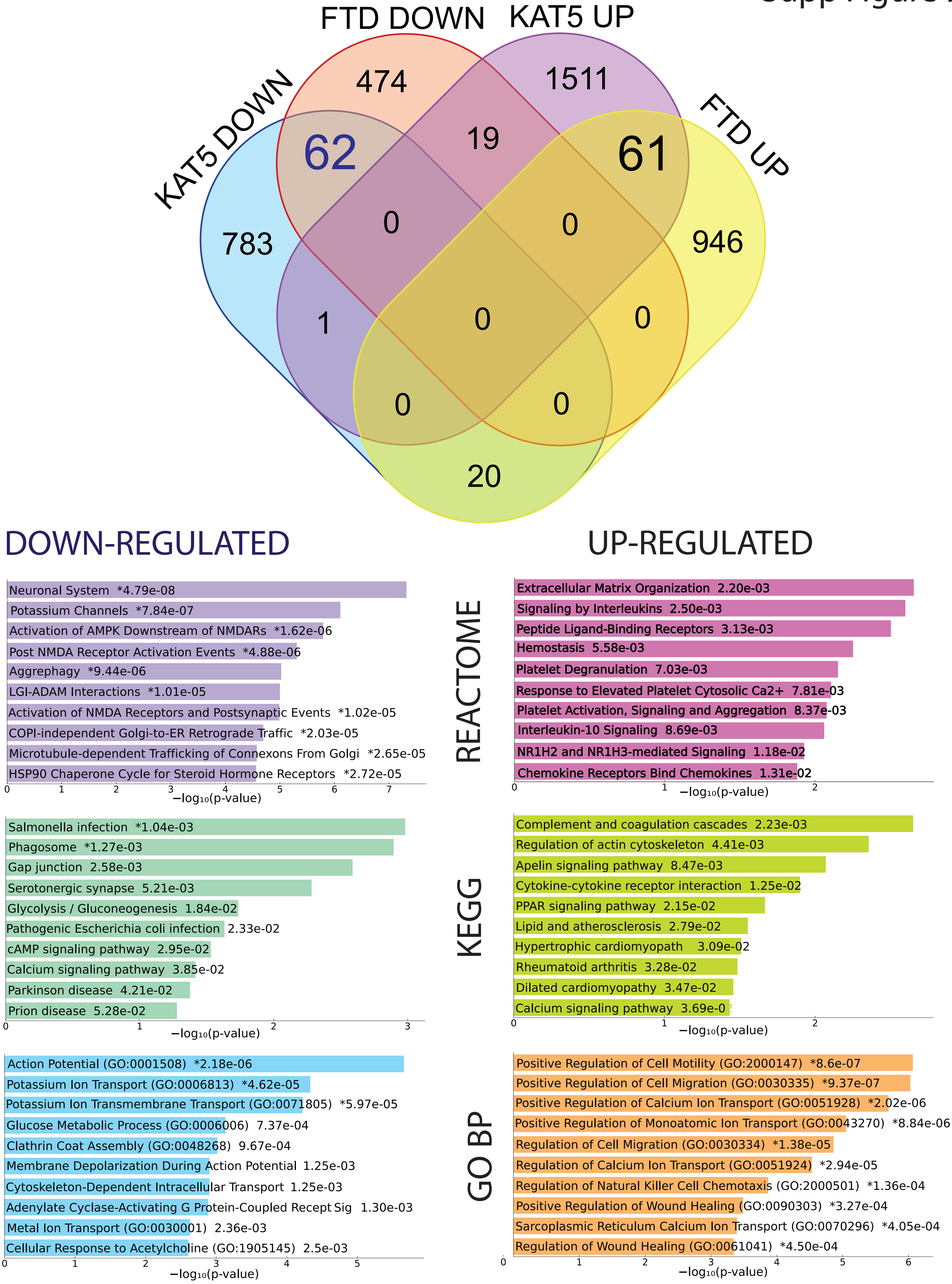

Figure Legend Supplemental 9. Kat5 cKO and MAPT FTD differential expression alignment.

The differential expression profile from the cKO Kat5 mouse were compared to the differential expression profile for the human frontotemporal dementia patients examined within the Risk and Modifying Factors in Frontotemporal Dementia (RIMOD-FTD) consortium. We specifically compared the MAPT linked FTD patients to the Kat5 cKO data to assess the most pathogenically similar dataset to AD. In all cases, there is 10-15% overlap in specific differentially expressed genes in up- and down-regulated gene sets. The degree of overlap is higher in the down-regulated FTD gene set. The overall biological pattern of enrichment in the up- and down-regulated gene set intersection is characterized by REACTOME and KEGG pathways in the top two plots (left: down-regulated; right: up-regulated). The GO biological process enrichment is shown on the bottom plots. The significance is corrected for multiple testing.
